# Differential S1PR Expression and Altered Barrier, and Inflammatory Marker Levels in Human Brain Endothelial Cells Following Acute Ischemia-Reperfusion Like Injury

**DOI:** 10.1101/2025.10.23.684217

**Authors:** Trevor S. Wendt, Nafis B. Eghrari, Matthew C. Lyons, Yasmin Beer, Rayna J. Gonzales

## Abstract

There are five known sphingosine-1-phosphate receptor (S1PR) types, and each type is coupled to diverse heterotrimeric G-protein subunits. S1PR type 1 (S1PR1) is highly expressed in the endothelium and receptor signaling pathways beneficially impact barrier function, thus there is a growing interest in the potential therapeutic role of S1PR1 during acute stroke injury. However, assessment of cerebrovascular S1PR1 and other S1PR expression profiles following acute ischemia-reperfusion injury have not yet been investigated. Therefore, we assessed the expression profile of human brain microvascular endothelial (HBMEC) S1PR1-5 following hypoxia plus glucose deprivation (HGD) exposure and HGD with reperfusion (HGD/R). We also investigated endothelial tight junction barrier and inflammatory mediator expression, indicative of an activated endothelial phenotype post HGD or HGD/R. Additionally, we investigated the effect of IL-1β, a downstream HGD induced inflammatory mediator, on endothelial mRNA expression of markers of function, barrier integrity, and inflammation and determined whether selective S1PR1 agonism altered mRNA expression of these genes. In terms of S1PR expression, a differential mRNA expression profile was observed revealing prominent basal endothelial expression of S1PR1-3. Following HGD, S1PR1 mRNA and protein expression increased with no observed changes in S1PR2 or S1PR3 protein levels. HGD/R increased endothelial inflammatory mediator and occludin mRNA expression with a temporal-dependent decrease in endothelial S1PR1 and claudin-5. Interestingly, we observed a positive and negative correlation between S1PR1 and S1PR2/3 with claudin-5, respectively. Selective S1PR1 agonism attenuated IL-1β induced MMP-9 mRNA and activity. In conclusion, HGD/R acutely increases endothelial cell expression of S1PR1 which presents a potential therapeutic target within the cerebrovasculature following ischemia-reperfusion injury. Additionally, HGD/R may potentially induce a phenotypic transformation within HBMECs, in which occludin expression predominates over claudin-5.

**NEW & NOTEWORTHY:** 

Graphical Abstract

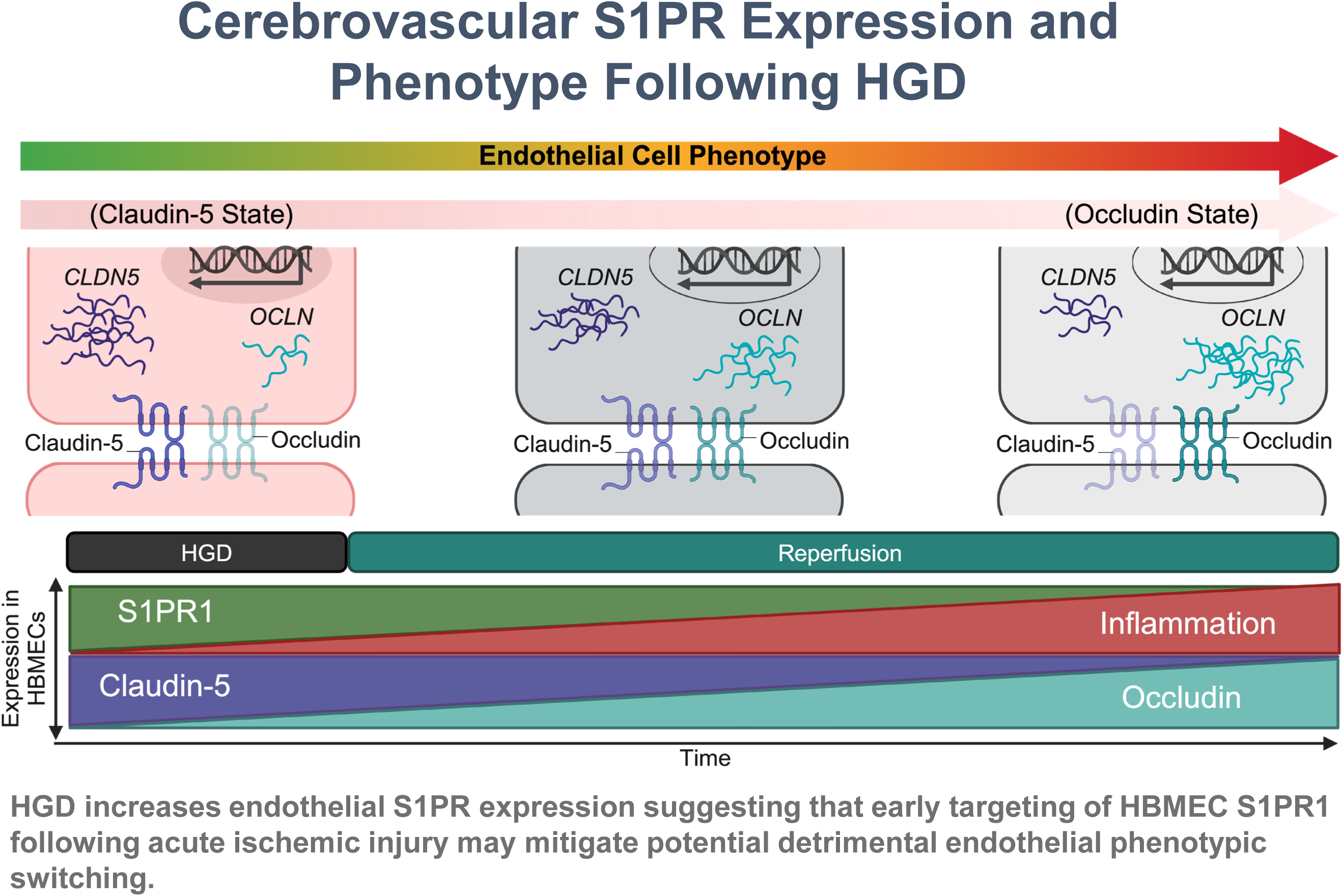

## 1. INTRODUCTION

Sphingosine-1-phosphate receptors (S1PRs) are part of the G-Protein Coupled Receptor (GPCR) family and play an important role in a diverse range of diseases and biological functions. Five receptor types have been reported to date and each is uniquely coupled with G-protein alpha subunits including Gαi, Gαq/11, and Gα12/13 to initiate multimodal downstream signaling pathways regulating cell migration, proliferation, growth, apoptosis, vascular integrity, and tone(1-3). The principal discovery of the *Edg-1* gene coding for S1PR1 was the first receptor type characterized within the GPCR family using human umbilical vein endothelial cells (HUVECs)(4). In the following decade, discovery of the remaining currently known S1PR genes coding for the S1PR2-5 types (5-8) were characterized and reported to be differentially expressed in various cell types within the heart, brain, immune, vasculature, and another organs systems in the body(9, 10). Further investigation into the expression profile of these S1PRs has been aimed towards understanding the complexity in regulating receptor activated signaling pathways linked to S1PRs during physiological and pathophysiological conditions.

It has been well established that the physiological responses elicited by S1PRs occur secondary to receptor activation by the endogenous ligand, sphingosine-1-phosphate (S1P)(11-15). Among the five receptor types, S1PR1/G_i_ activation is linked to T cell trafficking via receptor desensitization and consequent immune cell egress(16). However, in terms of the vascular endothelium, S1PR1 activation promotes endothelial barrier integrity preservation (17-20) and elicits vasodilation, primarily by inducing nitric oxide (NO) production via endothelial nitric oxide synthase (eNOS) activation(21, 22). Interestingly, both S1PR2 and S1PR3 are coupled to the G_i_, G_q_ or G_12/13_ alpha protein subunits [Reviewed in(23)] and the difference in their expression level determines unique downstream molecular consequences. Activation of either of these receptor types can modulate vasomotor tone and both have been shown to elicit either relaxation/dilation (24, 25) or constriction (26-30) depending on their vascular bed receptor type expression profile. For example, in both *in vivo* and *ex vivo* models, S1PR2 deletion blunted the vasoconstrictive response in mice induced by alpha-adrenergic activation and by exogenous application of S1P(28, 29). In contrast, using isolated isometric myography one study reported that S1P induced contraction in basilar artery rings from S1PR2 knockout mice compared to wildtype(26). S1PR3 also plays a bidirectional role in vascular tone regulation. In pre-contracted thoracic arteries, S1PR3 knockout attenuated the S1P-induced relaxation in mice(24). Conversely, S1PR3 activation promoted vasoconstriction in murine isolated cerebral arteries(30). S1PR3 has also been linked to vasculogenesis and angiogenesis by stimulating endothelial cell migration, similar to S1PR1(31).

As it relates to the human cerebrovasculature, the expression profile of S1PRs has not been well characterized, both basally and following acute ischemic injury. In this study, we aimed to delineate the expression profiles of S1PRs as well as several barrier molecules and inflammatory mediators in HBMEC exposed to hypoxia plus glucose deprivation (HGD), HGD with reperfusion (HGD/R; an *in vitro* ischemia-reperfusion injury), or treated with an HGD induced inflammatory mediator, IL-1β(32-35). Although S1PRs have been explored in the context of many different categories of diseases, including cancer, neurodegenerative diseases, and cardiovascular diseases, the mechanisms associated with S1PR activation in the pathological context of acute ischemic stroke (AIS) are still under investigation. The complex and varying effects of S1PR activation in the various vasculatures implicates the important role these receptors may play on brain blood vessel wall function and integrity during physiological and pathophysiological conditions and underline the gaining traction as potential therapeutic targets for AIS treatment.

The pathophysiology of AIS is complex, involving inflammatory pathways, oxidative stress, endothelial barrier disruption, excitotoxicity, apoptosis, angiogenesis, ionic imbalances, and neuropathology (36) of which cerebrovascular dysfunction is a key factor. Additionally, endothelial as well as vascular smooth muscle function are disrupted and have been characterized in part by alterations in endothelial tight junction proteins and vascular patency loss characterized by phenotypic switching of vascular smooth muscle to a synthetic phenotype(37, 38). The link between S1P/S1PR signaling and the pathophysiology of stroke is becoming more evident, particularly in the regulation of the BBB and endothelial function. A study using an *in vivo* middle cerebral artery occlusion (MCAO) mouse model reported increased mRNA levels of all five S1PR types in whole brain following ischemia and reperfusion(39). Others have demonstrated a beneficial role for S1PR1 on endothelial barrier integrity and reducing cerebral edema following stroke(40). However, not all studies provide a supportive role for S1PRs post stroke. There are studies that suggest that S1PR2 has been linked to endothelial activation and decreased barrier function(41, 42). In a pre-clinical ischemic stroke mouse model, S1PR2 activation led to cerebrovascular barrier disruption(43). Therefore, delineating the expression profile of S1PRs in human cerebrovascular cells merits further investigation to determine which receptor type(s) may be available for pharmacologic targeting during diseased states. Here we addressed the expression profiles of S1PRs in HBMECs as well as human brain vascular smooth muscle cells (**Supplemental Data**), both under basal conditions and following ischemic like conditions. We also addressed whether treatment with a selective S1PR1 agonist alters markers of endothelial function, barrier integrity, and inflammation.

## 2. MATERIALS AND METHODS

### 2.1. Cell Culture Model

Vendor-purchased cryopreserved subcultures of primary male human brain microvascular endothelial cells (HBMEC) (Cell Systems, Cat. No. ACBRI376, Lot No. 376.02.03.01.2F) and primary human brain vascular smooth muscle cells (HBVSMCs) (Cell Biologics, Cat. No. H-6085, Lot No. F11062014/17) were cultured as previously described(44, 45). In brief, HBVSMCs were grown in phenol red-free basal medium (Corning, Cat. No. 16-405-CV) supplemented with a smooth muscle cell growth supplement (insulin, human epidermal growth factor, human fibroblastic growth factor), gentamycin/amphotericin, and 5% FBS (Lonza; Cat. No. M1-3182). HBMECs were grown in phenol red-free Complete Classic Media (Cell Systems; Cat. No. 4Z0-500) containing 10% FBS (fetal bovine serum) and HBMECs were supplemented with Bac-off Antibiotic (Cell Systems; Cat. No. 4Z0-643) and CultureBoost (Cell Systems; Cat. No. 4CB-500). Cells reached 70%-80% confluence within 5-6 days and were then continued in culture and studied at Passage 7.

### 2.2. Hypoxia Plus Glucose Deprivation (HGD) and Reperfusion (HGD/R) Exposure

Cells were exposed to either normoxia or HGD conditions for 3h to mimic an in vitro acute ischemic injury. Immediately prior to hypoxic exposure, the media in the HBMEC and HBVSMC plates were replaced with modified DMEM (ThermoFisher Scientific; Cat. No. A14430-01) media containing no L-glutamine, phenol red, sodium pyruvate, and D-glucose. Plates were immediately placed in a humidified chamber (BioSpherix®) housed within a 5% CO_2_ incubator at 37°C and 1% O_2_ at room atmosphere for 3h following media replacement. Cells exposed to normoxic conditions were placed in a separate incubator at 5% CO_2_ and 21% O_2_. The hypoxic humidified chamber was filled with a medical grade gas mixture of 1% O_2_, 5% CO_2_, and nitrogen balance. HBMEC plates designated for HGD/R exposure had the modified DMEM medium replaced with Complete Classic Medium and were returned to 21% O_2_. Designated plates for temporal studies were either collected immediately following HGD 3h exposure or received fresh Complete Classic Medium and were then incubated in a CO_2_ incubator at 37°C at 21% O_2_ for 12h and 24h.

### 2.3. Recombinant IL-1*β* Exposure and Chemical Hypoxia

Recombinant IL-1*β* (Sigma; Cat No. I2393) as well as cobalt chloride (CoCl_2_; Sigma; Cat No. 232696) was utilized to interrogate specific intracellular mechanisms by which HGD/R may be altering markers of HBMEC function (eNOS), barrier integrity (CLDN5, OCLN, TJP1), and inflammation (ICAM1, OLR1, IL-1*β*, MMP9, and TIMP1). Recombinant IL-1*β* was prepared fresh under sterile conditions on the same day of administration for the planned experiments. Briefly, stock solutions (5*μ*g/mL) were made in ddH_2_O and further diluted in 1X Dulbecco’s phosphate buffer solution (DPBS) (Corning; Cat. No. 21-031-CV) to a final working concentration (400pg/mL). It has been previously reported that 5000pg/mL of IL-1*β* exposure temporally induced human cerebrovascular endothelial inflammation, activation, and barrier integrity loss(46). However, IL-1*β* concentrations have been found to increase in the serum to ∼80pg/mL in AIS patients within 24h of stroke onset(47). Therefore, to increase the translation of our observations we utilized a final dose of 400pg/mL for IL-1*β* exposure. We also utilized chemical induced hypoxia with CoCl_2_ to isolate the contributions of hypoxia on mediating changes to the cerebrovascular endothelium as it has been previously shown that CoCl_2_ disrupts the endothelial barrier(48). CoCl_2_ was prepared fresh under sterile conditions on the same day of administration for the planned experiments. Briefly, stock solutions (1,000mM) were made in ddH_2_O and further diluted in 1X Dulbecco’s phosphate buffer solution (DPBS) (Corning; Cat. No. 21-031-CV) to a final working concentration (100*μ*M).

### 2.4. Selective S1PR1 Ligand Treatment

Ozanimod (RPC1063, 5-[3-[(1S)-2,3-dihydro-1-[(2-hydroxyethyl)amino]-1H-inden-4-yl]-1,2,4-oxadiazol-5-yl]-2-(1-methylethoxy)-benzonitrile; S1PR1 agonist) (Cayman Chemical; Cat. No. 19922) was prepared fresh under sterile conditions on the same day of administration for planned experiments as previously described. (44, 49)Briefly, stock solutions (4mM) were made in dimethyl sulfoxide (DMSO) and further diluted in 1X Dulbecco’s phosphate buffer solution (DPBS) (Corning; Cat. No. 21-031-CV) to a final working concentration (0.5nM).

### 2.5. RNA Extraction and cDNA Synthesis

S1PRs 1-5 mRNA expression was assessed using reverse transcription polymerase chain reaction (RT-PCR) and quantitative real-time PCR (qRT-PCR). Briefly, cell pellets for each sample were collected using trypsin (ThermoFisher Scientific) and trypsin neutralizer (ThermoFisher Scientific) and centrifuged (Thermo Sorvall Legend RT+) at 3000 RPM for 10 minutes at 4°C. Steps were followed per the manufacturer’s instructions for RNA extraction (Qiagen kit; ThermoFisher Scientific, Cat. No. 74034). Pellets underwent a series of washes with the recommended buffers and centrifugations. The RNA was subsequently collected in 14ul of diethyl pryocarbonate (DEPC)-treated water. Purity and concentration of extracted RNA was confirmed using Nanodrop 2000 (ThermoFisher Scientific). RNA was then reverse transcribed to synthesize cDNA using the SuperScript™ III First-Strand Synthesis System (ThermoFisher Scientific; Cat. No. 18080051) according to manufacturer’s instructions. The resultant cDNA was used in RT-PCR or quantitative real-time RT-PCR reactions.

### 2.6. Reverse Transcription Polymerase Chain Reaction (RT-PCR)

RT-PCR was performed as described previously(44). In brief, reaction mixtures were prepared by loading equal amounts of cDNA determined by Nanodrop 2000, with 0.25μl of dNTP (10 μM) (ThermoFisher Scientific, Cat. No. 18427088), 1μl of 10X Taq Buffer (ThermoFisher Scientific, Cat. No. EP0702), 0.25μl of Taq polymerase (ThermoFisher Scientific, Cat. No. EP0702), 9.25μl of DEPC, 0.25μl of forward primer (10 μM) and 0.25μl of reverse primer (10 μM). Primers were designed in house using the NCBI and UCSC genome libraries and were obtained from Integrated DNA Technologies at 25nM scale with standard desalting. The RT-PCR protocol involved an initial melting step at 95°C for 3 min followed by 35 cycles of 95°C (denature) for 30 s, primer-specific annealing temperature for 30 s, and 72°C (elongation) for 60 s, then a 5 min incubation at 72°C in order to terminate the reaction. Primer annealing temperatures for S1PR1, S1PR2, S1PR3, S1PR4, S1PR5, and GAPDH are 59.6°C, 57.8°C, 59.6°C, 56.0°C, 58.1°C, and 57.8°C, respectively. The primer sequences used can be found in **Table 1**.

Following the PCR reaction samples were loaded alongside a UV standard (Bioland Scientific, Cat. No. DM01-01) onto a 2% agarose gel. Gels were run in a 1X tris-acetate-EDTA (TAE) (ThermoFisher Scientific, Cat. No. B49) solution for 15 min at 75V followed by 35 min at 95V. Gels were visualized and imaged using a ChemiDoc XRS UV gel imager to detect the gene of interest.

### 2.7. Quantitative Real-Time Polymerase Chain Reaction (qRT-PCR)

Reverse transcription qRT-PCR was carried out to assess S1PR1-3, CLDN5, OCLN, TJP1, PTGS2, IL-1β, and ICAM1 mRNA expression. RNA extraction and cDNA synthesis were performed as described in the Polymerase Chain Reaction section. Primer efficiencies were assessed via serial dilution from 100ng/μL to 0.01ng/μL and were found to be within 90-112% efficient. Following quantification (via Nanodrop 2000), cDNA from each sample was diluted to a working solution of 30ng/μL in DEPC. Duplicate or triplicate wells were used for each target within the 96-well plate (USA Scientific; Catalogue number: 1402-9100). Each well contained a mixture of cDNA (5μL), target primers diluted in DEPC (5μL; 0.4μM forward primer, 0.4μM reverse primer), and POWER SYBR Green PCR Master Mix (ThermoFisher Scientific; Catalogue number: 4368706). The 96-well plates were then centrifuged (Thermo Sorvall Legend RT+) at 1,000g for 2 min and then immediately loaded into the QuantStudio 6 and 7 Flex real-time PCR system (ThermoFisher Scientific). A two-step cycling protocol was utilized to collect cycle threshold (Ct) values of specific target amplicons and housekeeping gene GAPDH using the ΔΔCt relative quantification setting within QuantStudio 6. The RT-qPCR reaction involved 2 min at 50°C, an initial denaturing step at 95°C for 10min followed by 40 cycles of 95°C (denature) for 15 s and primer-specific annealing temperature for 1min. The primer annealing temperatures for S1PR1, S1PR2, S1PR3, CLDN5, OCLN, TJP1, PTGS2, IL-1β, ICAM1 and GAPDH were 65.5 °C. The primer sequences used can be found in **Table 2**.

### 2.8. Immunocytochemistry

Immunocytochemical analysis was used to assess S1PR1 and S1PR2 protein levels. In brief, HBMECs at density of 3 to 4 × 10^5^/cm^2^ were seeded onto glass coverslips coated with attachment factor (Cell Systems; Catalogue number: 4Z0-201) within 6-well plates at passage 7, grown to 70-80% confluency, and divided into two groups as described before: normoxia and HGD. Following 3h exposure, plates were immediately removed from the chamber and placed on ice, media was aspirated, and cells were washed once with ice-cold DPBS. HBMECs were then fixed with 4% formaldehyde (10 min, RT) and subsequently washed twice for 4 min with DPBS. HBMECs were then permeabilized with 0.1% TritonX-100 (Catalogue number: T9284) for 10 min and then blocked in 2% BSA (15 min, RT). S1PR1 expression and nuclear localization was determined using anti-S1PR1 (Invitrogen; Catalogue number: PA1-1040) and S1PR2 expression using anti-S1PR2 (Santa Cruz Bio; Catalogue number: sc-365589) at concentrations 1:500 and 1:100, respectively, in 2% BSA. Following an overnight incubation at 4°C, HBMECs were washed by 2% BSA (3×, 5 min) followed by secondary antibody incubation with either Alexa Fluor 488 goat anti-mouse (ThermoFisher Scientific; Catalogue number: A11001) or Alexa Fluor 555 goat anti-rabbit (Thermo Fisher Scientific; Cat. No. A21428) at 1:6,000 in 2% BSA, for 1h at RT protected from light. A quick wash was then performed with the 2% BSA following by washes with DPBS (3×, 5 min). Two coverslips from each treatment group were incubated in secondary antibody only and served as negative controls. Next, excess DPBS was removed, and the coverslips were mounted onto slides utilizing vectashield mounting medium with DAPI (Vector Laboratories; Catalogue number: H-1200). Coverslips were then sealed onto slides using nail polish and images captured with an inverted digital microscope (Keyence; Catalogue number: BZ-X800) followed by area quantification using ImageJ software. This area quantification is described in detail in our previous studies in both the human brain vascular smooth muscle and microvascular endothelium(44, 45).

### 2.9 In-Cell Western Blot

Protein levels for iNOS and claudin-5 were examined following HGD/R exposure as previously described(44). In brief, in-cell western blotting was accomplished via CellTag 520 LiCor Stain Kit IV (LiCor Biosciences; Cat. No. 926-42094) according to the manufacturer’s instructions. HBMECs at density of 1 × 10^5^/cm^2^ were seeded into 96-well plates (Greiner Bio-One; Cat. No. 655097) at *passage 7* and grown to 85-90% confluency. Following 3h/24h exposure/treatment, plates were immediately removed from incubator/sub chamber, media aspirated, and cells fixed with 3.7% formaldehyde for 20min at room temperature. Following fixation, cells were incubated with blocking solution Intercept (TBS) Blocking Buffer (LiCor Biosciences; Cat. No. 927-60001) for 1.5h with gentle agitation at room temperature. The cells were then probed overnight at 4°C with iNOS (Cell Signaling; Cat. No. 9543) or claudin-5 (ThermoFisher Scientific; Cat. No. 34-1600) primary antibodies which were all diluted to 1:500 in TBST. After TBST wash cycle (5×, 5min) with gentle agitation at room temperature, cells were incubated in goat anti-rabbit IRDye 800CW dye (LiCor Biosciences) or goat anti-mouse IRDye 680CW dye (LiCor Biosciences) secondary antibodies both conjugated with CellTag 520 Stain (LiCor Biosciences) at room temperature for 1h. Following another 5×, 5min TBST wash cycle, protein expression was visualized using an Odyssey Classic infrared imager (LiCor Biosciences). Protein fluorescent densities were then analyzed using ImageStudio 3.0 software (LiCor Biosciences) and normalized to cells stained by CellTag 520 which served as a loading control.

### 2.10. Zymography

Zymography experiments were conducted as previously described(44). Briefly, media was extracted from cultures following HGD exposure/drug treatment. The media was briefly centrifuged at 1,000g for 5min to remove any cellular debris and the supernatant was removed utilized as conditioned media. Total protein within conditioned media was determined on the day of zymography experimentation and determined by using a bicinchoninic acid protein assay kit (ThermoFisher Scientific) according to manufacturer’s protocol and measured on a Safire II (Tecan) plate reader. Conditioned media samples were diluted in Tris-glycine SDS sample buffer and the equally diluted samples and fluorescent standards (LI-COR Biosciences) were loaded onto Novex 10% zymogram plus gelatin protein gels (ThermoFisher Scientific; Cat. No. ZY00100BOX). Protein separation occurred by SDS-PAGE using a mini–PROTEAN Tetra electrophoresis system (Bio-Rad Laboratories) at 100V for 2h and held at 4°C. After electrophoresis, the gel was incubated in 1× zymogram renaturing buffer (ThermoFisher Scientific; Cat. No. LC2670) for 0.5h at RT with gentle agitation. Zymogram renaturing buffer was decanted and replaced with 1× zymogram developing buffer (ThermoFisher Scientific; Cat. No. LC2671) and allowed to equilibrate the gel for 0.5h at RT with gentle agitation. Following equilibration, developing buffer was decanted and gels washed with nano-pure H_2_O and replaced with fresh 1× zymogram developing buffer and developed for 24h at 37°C. Following developing, gels were briefly washed with nano-pure H_2_O and stained with Coomassie blue staining solution (2.5g Coomassie Brilliant blue R-250, 450ml 100% methanol, 100ml 100% acetic acid, 400ml nano-pure H_2_O) for 2h at RT with gentle agitation. Gels were then washed with nano-pure H_2_O and de-stained with Coomassie blue de-staining solution (450ml 100% methanol, 100ml 100% acetic acid, 400ml nano-pure H_2_O) for 40min at RT with gentle agitation. The area of gelatin degradation at approximately 82kDa and 63kDa was indicative of MMP-9 and MMP-2 activity respectively and appeared as distinct bands. The intensities of both MMP-9 and MMP-2 were obtained by an Odyssey Classic infrared imager (LI-COR) and inverse bands densities were then analyzed using ImageStudio 3.0 software (LI-COR).

### 2.11. Reagents

All reagents and chemicals were obtained from Sigma Aldrich (St Louis, MO) unless otherwise noted.

### 2.12. Data and Statistical Analysis

For western blot analyses, each treatment group was run on the same gel to enable direct comparison. Data were expressed as fluorescent signal density of the desired protein normalized to the fluorescent signal density of β-actin (loading control) and fold changed to the average of the normalized normoxic values. mRNA data generated from qPCR experimentation were expressed as either relative expression (2^-^Δ^ct^) normalized to GAPDH (loading control) or as normalized to normoxic values (2^-^ΔΔ^ct^) normalized to GAPDH. Experiments were repeated for statistical analysis and data graphed using Prism 9.3.0 for Windows, GraphPad Software, USA, www.graphpad.com (GraphPad Software). Confirmation of normal distribution was achieved with a Shapiro-Wilk test. F-test confirmed if data sets achieved equal variance among groups. For analysis of datasets that achieved a normal distribution, direct comparisons between two groups were made using an unpaired t test alone if equal variance was achieved or with Welch’s correction if equal variance was not achieved. Comparisons between three or more groups were made using a repeated one-way ANOVA with Tukey’s multiple comparisons post hoc test indicated in the respective figure legends. Datasets that did not achieve a normal distribution underwent direct comparisons via Mann-Whitney or multiple comparisons via Kruskal–Wallis tests. Simple linear regression was utilized to assess relationship between S1PR1-3 and associated barrier and inflammation markers. P < 0.05 was considered significant. Values are expressed as means ± SD.

## 3. RESULTS

### 3.1. HBMEC S1PR mRNA expression profile following in vitro HGD exposure

The vascular endothelium is known to highly express S1PR1 with comparatively lower reported expression levels of S1PR2 and S1PR3(50-52), however a comprehensive expression profile of S1PRs in the human brain vasculature has not been addressed. Thus, we assessed S1PR mRNA expression profiles in primary cultured HBMECs (**Fig. 1**) and HBVSMCs (**Supplemental Fig. 1**). Based on RT-PCR and agarose gel electrophoresis, S1PR types 1-4 mRNA expression was detected in HBMECs (**Fig. 1A**). Semi-quantitative analysis of these receptors relative to GAPDH revealed expression of S1PR1-3 with very low levels of S1PR4 and no detectable amplicons for S1PR5. HBVSMCs exhibited a similar S1PR1-4 mRNA expression profile however an additional product corresponding to S1PR5 appearing as a faint band at 108bp was also visualized (**Supplemental Fig. 1A**). Since S1PR1-3 amplicon products were the most prominent product detected in our vascular cell models, we next focused on assessing the impact of HGD, an in vitro ischemia-like stroke injury model, on types 1-3 mRNA expression levels via qRT-PCR (**Fig. 1B**). We observed that HBMEC S1PR1 mRNA significantly increased following 3h of HGD exposure. In contrast, HGD induced a significant decrease in S1PR1 mRNA expression in HBVSMCs (**Supplemental Fig. 1B**). Furthermore, HGD exposure had no effect on S1PR2 mRNA expression levels in either HBMECs or HBVSMCs. In terms of S1PR3, following HGD exposure, S1PR3 mRNA was increased in HBMECs and decreased in HBVSMC. Together these data suggest that HGD 3h exposure may differentially alter HBMEC and HBVSMC S1PR3 at the transcriptional level in a manner similar to S1PR1. Previous studies have demonstrated that S1PR mRNA quantification of HUVECs resulted in S1PR1 > S1PR3 > S1PR2(53, 54). In line with these studies, using qRT-PCR we confirmed that HBMEC mRNA levels rank S1PR1>>S1PR3>S1PR2 both basally and following HGD (**Fig. 1C**). With regards to vascular smooth muscle, S1PR1 has been reported to be the predominant receptor studied(55, 56). In a previous study using HVSMCs (origin of vascular bed not disclosed), the S1PR expression profile was similar to what we observed in our HBVSMC model(55). In our studies, we noted that unlike HBMECs, under normoxic and HGD conditions HBVSMC the expression profile resulted in S1PR3>S1PR2>>S1PR1 (**Supplemental Fig. 1C**). Based on these findings, the contrasting relative mRNA expression of S1PRs in our two cell types of interest may suggest a differential bioavailability of these receptors for binding by the endogenous S1P ligand and subsequential targeting by pharmacological ligands.

**Figure 1.**
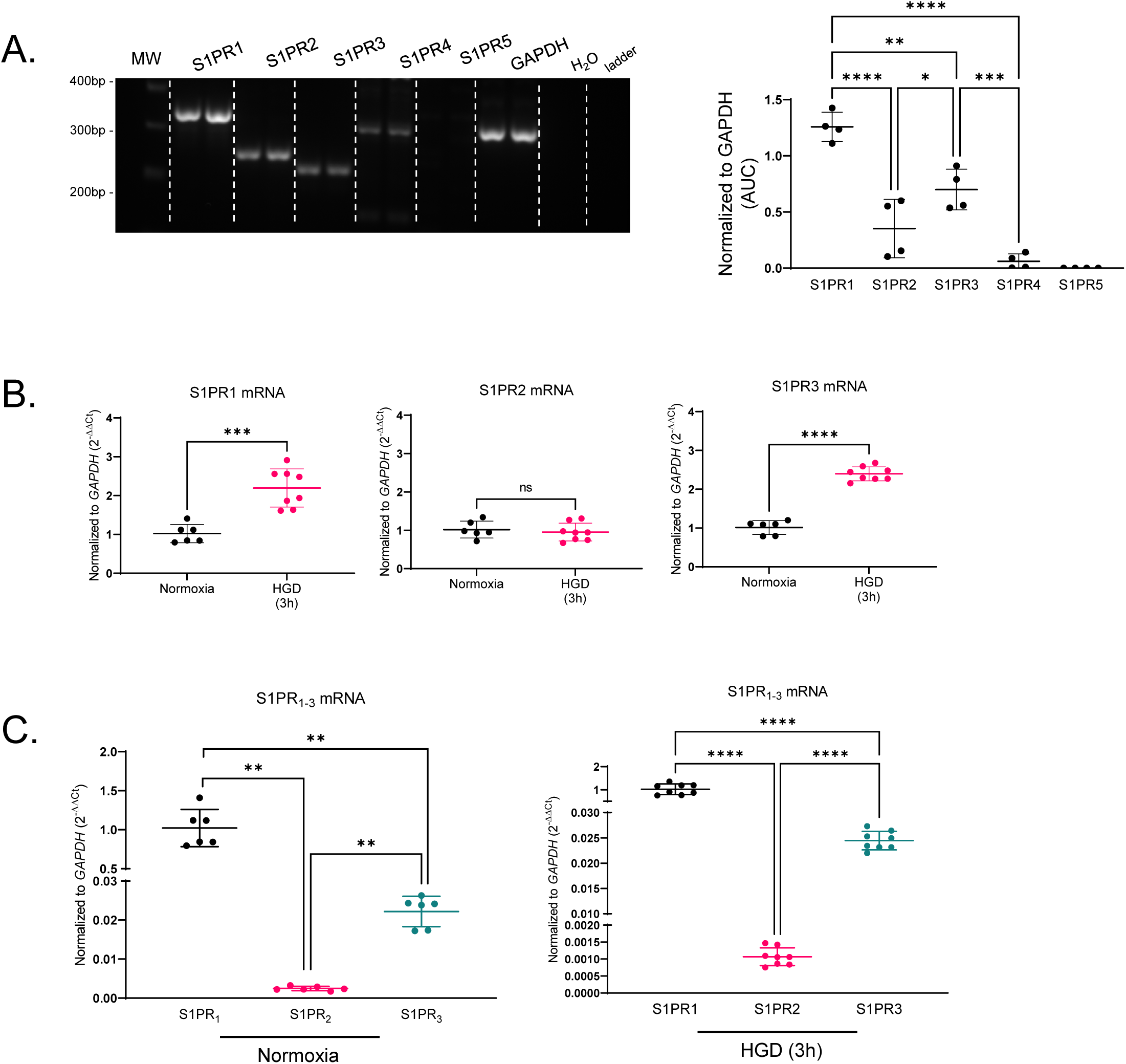
Relative S1PR Types 1-5 mRNA Expression Profiles in Cultured Human Brain Endothelial Cells Following Normoxia or HGD. (**A**) RT-PCR ran on 2% agarose gels for sphingosine-1-phosphate receptor (S1PR) types 1-5 to detect receptor mRNA presence in human brain microvascular endothelial cells (HBMEC). GAPDH (loading control) yielded a product size of 309 bp. S1PR1 yielded a product size of 343 bp, S1PR2 yielded a product size of 246 bp, S1PR3 yielded a product size of 215 bp, S1PR4 yielded a product size of 200, and S1PR5 yielded a product size of 104 bp. Bar graph illustrates semi-quantitative densiometric analysis of detected S1PR transcription normalized to GAPDH. n=4. (**B**) Bar graphs depict qRT-PCR mediated quantitation of S1PR1, S1PR2, and S1PR3 mRNA present within HBMECs exposed to either normoxia or HGD (3h) normalized to the housekeeping gene (GAPDH) and expressed as 2^−ΔΔCt^. n=6-8. (**C**) qRT-PCR graphs of relative S1PR1, S1PR2, and S1PR3 mRNA expression in HBMECs following normoxia and 3h HGD exposure normalized to the housekeeping gene (GAPDH) and expressed as 2^−ΔΔCt^. n=6-8. Direct comparisons were performed with unpaired t test with Welch’s correction. Multiple comparisons were performed with one-way ANOVA with Tukey’s post-hoc tests. *P<0.05, **P<0.01, ***P<0.001, ****P<0.0001.

### 3.2. HGD elicited a selective increase in S1PR1 protein levels without altering S1PR2-3 levels in HBMECs

After examining the S1PR mRNA expression profile in HBMECs, we next assessed whether the protein levels of S1PR1-3 would be reflective of the previous mRNA findings and if HGD would affect protein expression (**Fig. 2**). Utilizing immunocytochemical quantification analysis, as described previously(44, 45), we observed a significant increase in S1PR1 protein levels following HGD 3h exposure (**Fig. 2A**). In terms of S1PR2, in line with our previous observations at the mRNA level, S1PR2 protein levels were not different in the normoxic vs. HGD groups (**Fig. 2B**). However, unlike the increase in S1PR3 mRNA expression, S1PR3 protein levels were not increased following 3h HGD (**Fig. 2C**). These data further support our prior observations that S1PR1 is upregulated acutely following an ischemic-like injury and could serve as a potential therapeutic target in mitigating endothelial pathogenesis during stroke.

**Figure 2.**
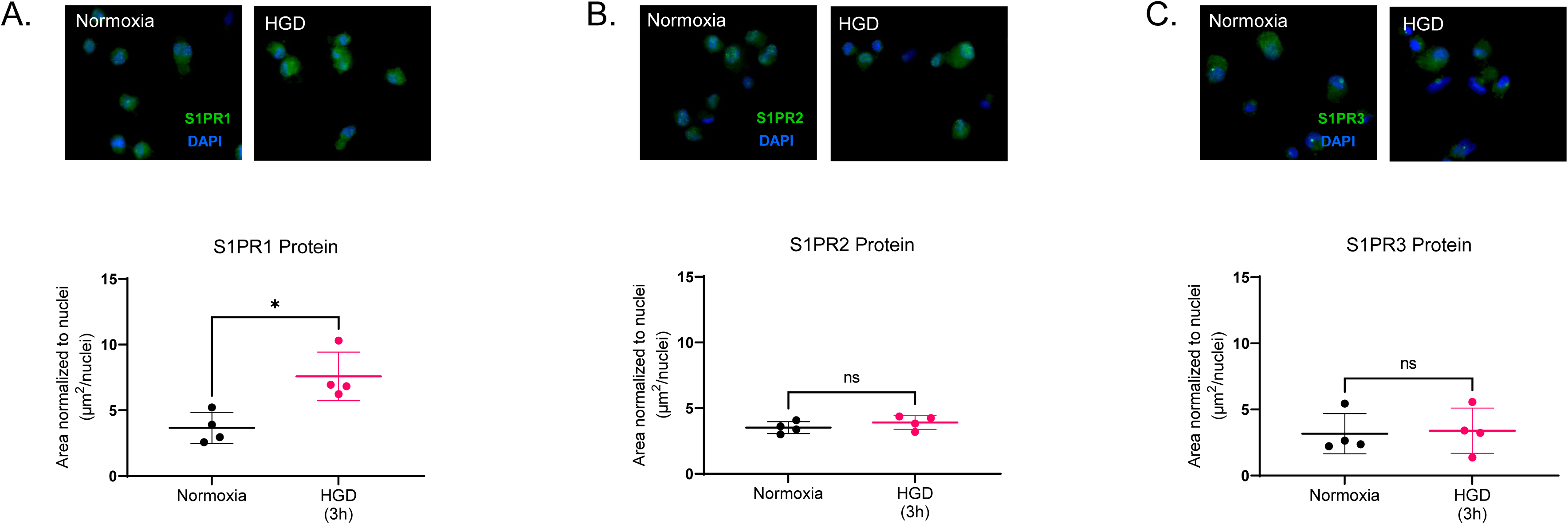
HGD-Mediated Alterations in HBMEC S1PR Types 1-3 Protein Levels. (**A-C**) Representative immunocytochemical images of anti-S1PR1 (S1PR1; green), anti-S1PR2 (S1PR2; green), and anti-S1PR3 (S1PR3; green) in HBMECs following normoxia or exposure to 3h HGD. Imaged at 40X magnification. Nucleus is labeled with 4’,6-diamidine-2’-phenylindole dihydrochloride (DAPI; blue). Scale bar, 50μM. Bar graphs depicts the relative quantification of (**A**) S1PR1, (**B**) S1PR2, (**C**) S1PR3 in HBMECs normalized to the number of nuclei and expressed as a ratio of the area occupied by S1PR1 to number of nuclei (μm^2^/nuclei). n=4. Direct comparisons were performed with unpaired t test with Welch’s correction. Mann-Whitney’s test was utilized as data are non-parametric. *P<0.05.

### 3.3. HGD-induced differential HBMEC barrier and inflammatory mRNA expression

Key components in the progression of cerebral endothelial pathogenesis during acute ischemic injury include aspects such as inflammation and tight junction disruption(57). In this set of experiments, we assessed mRNA levels of the pro-inflammatory mediators COX-2 and IL-β as well as adhesion molecule ICAM1 following HGD which have all been implicated in contributing to the pathogenesis of acute ischemic injury(58-61). We also assessed claudin-5, occludin, and zonula occludens 1 (ZO-1) as they have been well characterized as integral to maintaining endothelial barrier integrity and are disrupted following ischemic stroke(62-66). Barrier and inflammatory mRNA expression levels were measured immediately following HGD 3h exposure (**Fig. 3**). We hypothesized that HGD would concomitantly increase pro-inflammatory and adhesion marker mRNA expression and decrease barrier integrity marker expression.

**Figure 3.**
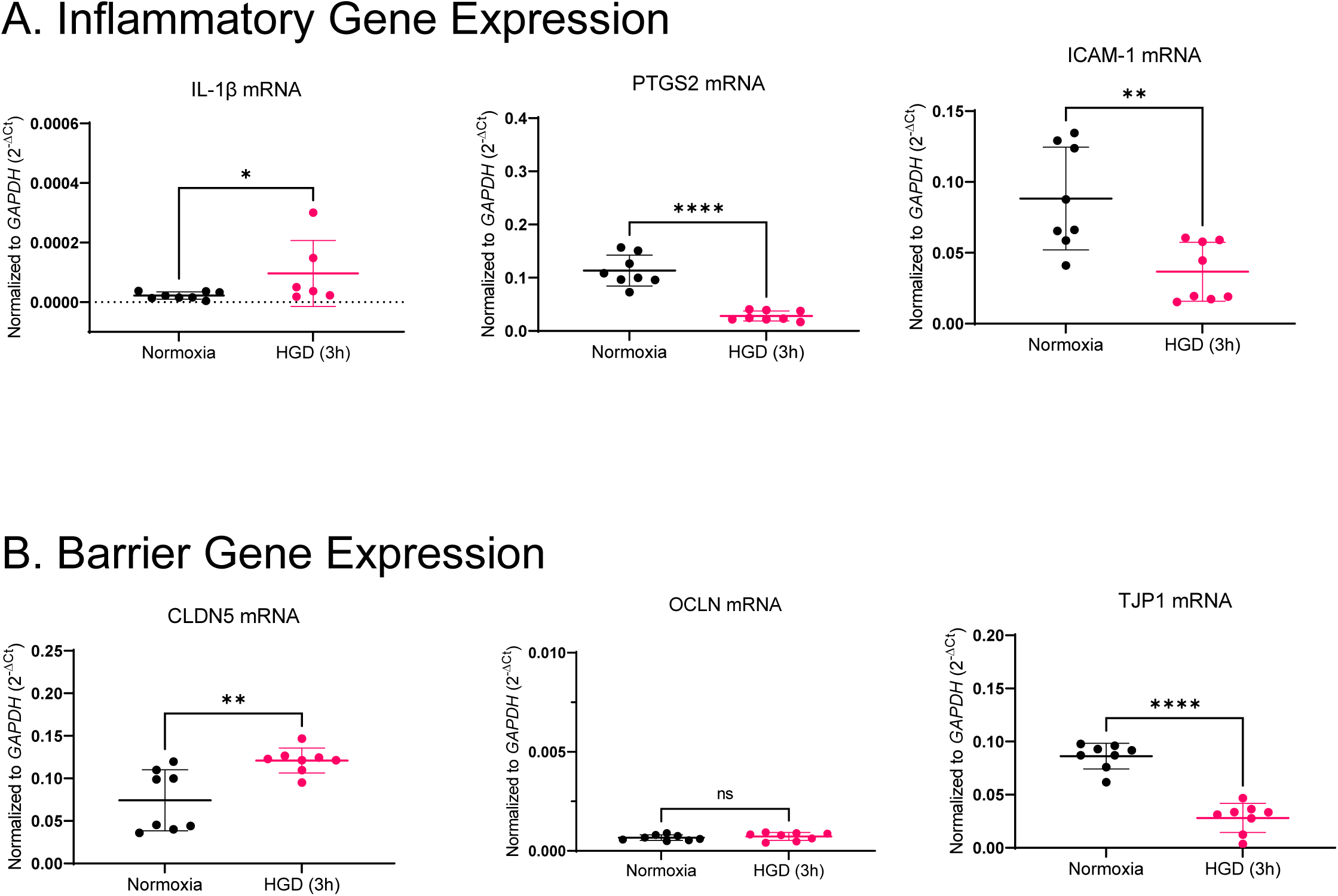
HBMEC Inflammatory and Barrier mRNA Expression Following HGD Exposure. qRT-PCR graphs of (**A**) *IL-1β, PTGS2*, *ICAM-1* as well as (**B**) *CLDN5*, *OCLN,* and *TJP1* mRNA in HBMECs following normoxia and HGD 3h exposure normalized to the housekeeping gene (GAPDH) and expressed as 2^−ΔCt^. n=6-8. Direct comparisons were performed with unpaired t test with Welch’s correction. Mann-Whitney’s test was utilized for analysis of *IL-1β, ICAM-1*, and *CLDN5* as these data are non-parametric. *P<0.05, **P<0.01, ****P<0.0001.

Consistent with our hypothesis, we observed a modest but significant increase in levels of IL-1β (**Fig. 3A**). Counter to the hypothesis, there was a notable HGD-induced decrease in PTGS2 and ICAM1 mRNA. In terms of barrier expression, claudin-5 mRNA expression increased following HGD 3h exposure with a concomitant decrease in ZO-1 expression (**Fig. 3B**). Interestingly however, while there was an increase in claudin-5 mRNA expression at this time point, there was no difference in occludin mRNA expression following 3h HGD exposure. Collectively, these findings indicate that acutely following a 3h in vitro ischemic-like injury, human brain endothelium express differential levels of pro-inflammatory mRNA expression profiles while concomitantly displaying similar a paradoxical barrier marker mRNA expression profile. These findings prompted us to further investigate the potential impact of simulated reperfusion injury on these outcomes.

### 3.4. S1PR1 mRNA expression and protein levels were concomitantly decreased following HGD 3h and with 24h reperfusion

Selective S1PR1 ligands/modulators including ozanimod (20-22 h)(67), amiselimod (386-423 h)(68), ponesimod (21.7 and 33.4 h)(69), and siponimod (56.6 h) (70)exhibit a range of half-life’s that may provide clinical relevance when studying endothelial S1PR1 as a target receptor during the potential therapeutic window in patients afflicted with AIS. Thus, we next assessed expression of S1PR1-3 following HGD 3h and 24h post-ischemic-like injury. With growing interest in S1PRs as promising therapeutic targets, understanding how the expression of these receptors evolves over time and during reperfusion remains to be fully understood. Several recent in vitro ischemia-reperfusion studies performed within lung epithelial cells(71), cardiomyocytes(72-74), and HUVECs (75, 76) along with our previous study in primary human brain microvascular endothelial cells (77) which have demonstrated the detrimental impact of reperfusion injury. Following HGD 3h with a 24 reperfusion (HGD 3h/R 24h) we detected a decrease in S1PR1 mRNA expression which was not observed with S1PR2 or S1PR3 (**Fig. 4A**). Since S1PR1 mRNA was decreased, we further interrogated protein levels and noted a similar decrease in protein levels (**Figs. 4B**). To gain an appreciation for the temporal change in expression profile of S1PR1-3 in response to the 24h reperfusion phase we also assessed S1PR1-3 mRNA expression in cells exposed to HGD 3h with 12h reperfusion (**Supplemental Fig. 2A**). No difference was noted in the mRNA expression of S1PR1 or S1PR2; and interestingly a decrease in S1PR3 mRNA was detected. Line graph in **Fig. 4C** summarizes of the relative temporal mRNA expression profile of S1PR1-3 following either normoxia or HGD 3h/R 0h, 12h, and 24h. Together, these data suggest that initially following acute HGD exposure, there is an increase in S1PR1 that gradually decreases during the simulated reperfusion phase. Our findings promote the notion that early therapeutic treatment targeting cerebral endothelial S1PR1 could potentially have the most robust response if given following an acute ischemic event and within the first 12h during reperfusion. Moreover, the relatively low expression and alteration in expression of both S1PR2 and S1PR3 compared to S1PR1 within the HBMECs further supports S1PR1 as a strong therapeutic target within the HBMECs following acute cerebral ischemic injury (**Supplemental Fig. 2B**).

**Figure 4.**
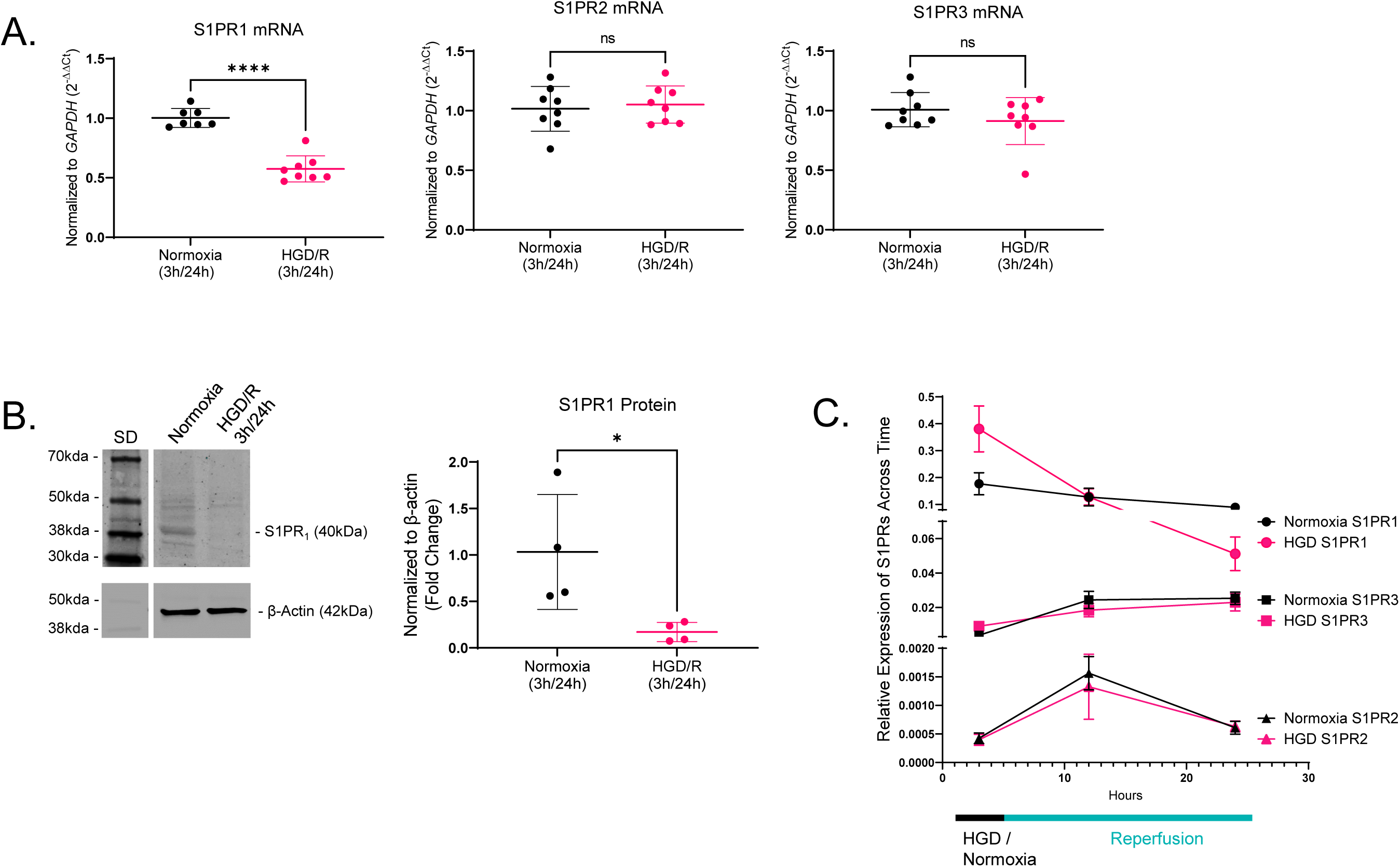
HBMEC S1PR Type 1-3 mRNA and Protein Expression Following In Vitro Hypoxia Plus Glucose Deprivation/Reperfusion (HGD/R). qRT-PCR graphs of (**A**) S1PR1, S1PR2, and S1PR3 mRNA expression in HBMECs following normoxia and HGD 3h /R 24h exposure normalized to the housekeeping gene (GAPDH) and expressed as 2^−ΔΔ*Ct*^. n=7-8. (**B**) Representative western blot demonstrating the band migration of S1PR1 (40kDa) and *β*-actin (42kDa; same blot stripped and re-probed) in HBMECs following exposure to normoxia or HGD 3h /R 24h. Graph illustrates densiometric analysis of S1PR1 protein levels normalized to *β*-actin and expressed as fold change to normoxia. n=4. (**C**) Graphical summary of the temporal relative transcript profile of HBMEC S1PRs 1-3 following either normoxia 3h or HGD 3h with reperfusion for 0h, 12h, and 24h. n=4-8. Direct comparisons were performed with unpaired t test with Welch’s correction. Mann-Whitney’s test was utilized for analysis of S1PR3 as data are non-parametric. *P<0.05, ****P<0.0001.

### 3.5. HGD followed by simulated reperfusion decreased HBMEC barrier markers and concomitantly increased inflammatory mediators

Recent advancements in techniques allowing for resolution of ischemia and establishment of reoxygenation has amplified the impact of reperfusion injury ictus including elevation of reactive oxygen species, inflammation, and BBB weakening [Reviewed in(78)]. Therefore, we next addressed the impact of HGD 3h/R 24h on inflammatory and endothelial barrier markers (**Fig. 5**). Counter to our findings conducted at HGD 3h in the absence of perfusion (**Fig. 3**) and HGD 3h/R 12h (**Supplemental Fig. 3A**), we noted an increase in IL-1β, PTGS2 mRNA expression as well as an increase in levels of the vascular inflammation marker, iNOS, following HGD 3h/R 24h exposure (**Fig. 5A-B**). Although following exposure to HGD alone we detected a decrease in ICAM-1, we did not observe any temporal differences in adhesion receptor mRNA expression following HGD 3h/ R 24h (**Fig. 5A**) or HGD 3h/R 12h (**Supplemental Fig. 3A**). Next, we assessed ischemic like injury with reperfusion altered endothelial barrier marker expression.

**Figure 5.**
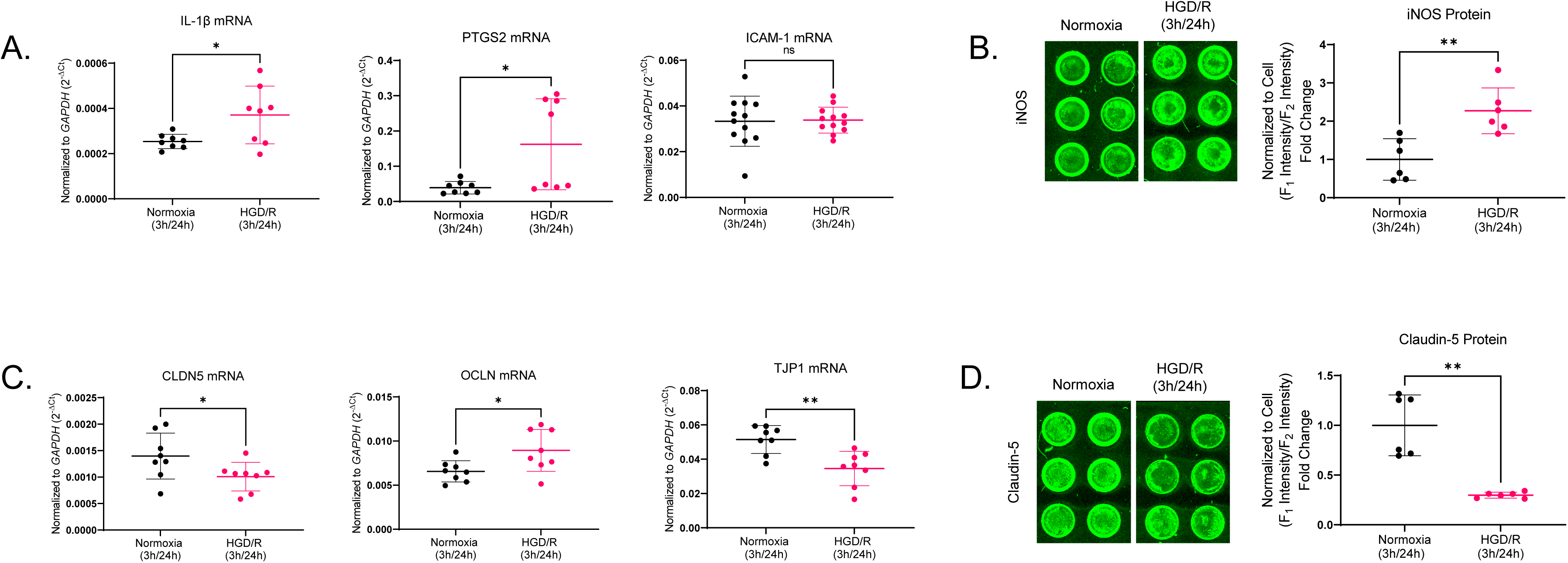
HBMEC Inflammatory and Barrier mRNA and Protein Expression Following Hypoxia Plus Glucose Deprivation/Reperfusion (HGD/R) Exposure. qRT-PCR graphs of (**A**) *IL-1β*, *PTGS2*, and *ICAM-1* mRNA in HBMECs following normoxia and HGD 3h /R 24h exposure normalized to the housekeeping gene (*GAPDH*) and expressed as 2^−ΔCt^. n=8. (**B**) Representative in cell western images of HBMECs exposed to normoxia or HGD 3h /R 24h. Graph depicts anti-iNOS normalized to cell labeling fluorescence, F1 and F2 respectively, by densiometric analysis of iNOS protein expression. n=6. qRT-PCR graphs of (**C**) *CLDN5*, *OCLN*, and *TJP1* mRNA expression in HBMECs following normoxia and HGD 3h /R 24h exposure normalized to the housekeeping gene (GAPDH) and expressed as 2^−ΔCt^. n=8. (**D**) Representative in cell western images of claudin-5 labeling in HBMECs exposed to normoxia 3h or HGD 3h with R 24h. Graph depicts anti-claudin-5 normalized to cell labeling fluorescence, F1 and F2 respectively, by densiometric analysis of claudin-5 protein expression. n=6. Direct comparisons were performed with unpaired t test with Welch’s correction. Mann-Whitney’s test was utilized for the analysis of *PTGS2* as data are non-parametric. *P<0.05, **P<0.01.

Following an HGD 3h /R 24h exposure we observed a decrease in TJP1 and CLDN5 mRNA expression with a paradoxical increase in OCLN mRNA expression (**Fig. 5C**). Similar to CLDN5 mRNA expression, HGD 3h /R 24h decreased claudin-5 protein levels suggesting potential barrier dysfunction. Together, these data suggest that there is a rise in brain endothelial derived inflammation, involving potential mediator including COX-2, IL-1*β*, and iNOS which is accompanied by a decrease in ZO-1 and claudin-5. Intriguingly however, there was a notable increase in occludin expression, potentially suggesting an endothelial phenotypic switch of preferential claudin-5 to occludin expression at 24h post-HGD injury. Further investigation into the functional impact of this potential phenotype expression of the endothelium is warranted but exceeds the scope of this study.

### 3.6. Temporal dependent alterations in barrier and inflammatory marker expression following simulated ischemic like injury with reperfusion

Here we assessed the temporal expression of brain endothelial barrier markers TJP1, CLDN5, and OCLN in HBMECs exposed to either normoxia or HGD 3h with reperfusion at 0, 12h, and 24h (**Fig. 6**). Thus far we have investigated the temporal impact of HGD/R exposure relative to the normoxic control groups. This experiment was designed to directly assess the relative mRNA expression of select barrier markers within each treatment condition (i.e., normoxia or HGD/R). We observed that under normoxic exposure at 3h/0h, HBMECs expressed similar mRNA levels of TJP1 and CLDN5 which were both significantly greater than occludin levels (**Fig. 6A**). As the cells continued to grow and reached the 3h/12h time point, we observed a decrease in CLDN5 mRNA levels relative to TJP1; however, CLDN5 mRNA expression levels remained significantly greater than OCLN. Finally, at the normoxic 3h/24h time point, HBMECs elicited greater OCLN levels relative to CLDN5, both of which were less than TJP1. Similarly, in HBMECs exposed to HGD/R we observed a relatively higher expression of TJP1 and CLDN5 compared to OCLN at 3h/0h (**Fig. 6B**). However, unlike the normoxic control HBMECs at 3h/0h, CLDN5 was most highly expressed following HGD 3h/R 0h exposure. Moreover, at 3h/12h there was a notable increase in TJP1 levels relative to both CLDN5 and OCLN which were not different from each other. Then at 3h/24h, similar to HBMECs exposed to normoxic control conditions, there was an increase in OCLN levels relative to CLDN5. Overall, these data suggest that both under normoxic and HGD/R conditions TJP1 mRNA expression remain relatively most highly expressed compared to CLDN5 and OCLN in HBMECs (**Fig. 6A-C**). Moreover, it appears that under HGD/R the potential phenotype switch from CLDN5 to OCLN may be induced sooner compared to normoxic controls. In an experiment where HGD was increased to 6h and included R 12h (HGD 6h/R 12h), the increased duration of HGD appeared to further exacerbate this increase in OCLN compared to HGD/R (3h/12h) (**Supplemental Fig. 3A**).

**Figure 6.**
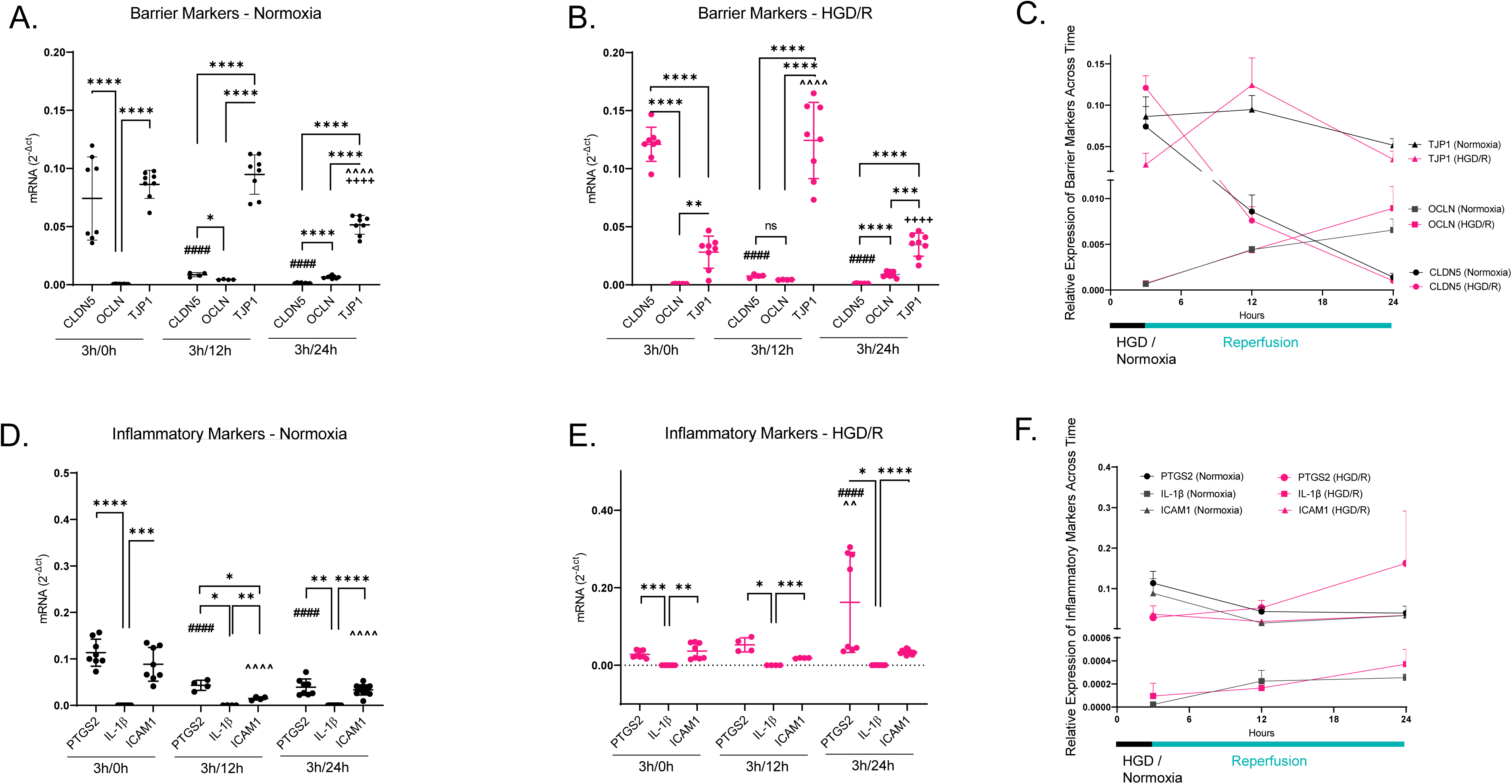
Temporal Barrier and Inflammatory Marker mRNA Expression Profile in HBMECs Following HGD/R Exposure. *CLDN5*, *OCLN*, and *TJP1* relative mRNA expression in HBMECs following (**A**) normoxia 3h or (**B**) HGD 3h followed by 0, 12, and 24h of simulated reperfusion normalized to the housekeeping gene (GAPDH) and expressed as 2^−Δ*Ct*^. n=4-8. (**C**) Graphical summary of the temporal relative expression profile of *CLDN5*, *OCLN*, and *TJP1* following either normoxia 3h or HGD 3h with reperfusion for 0, 12, and 24h. n=4-8. *PTGS2*, IL-1β and *ICAM1* relative mRNA expression in HBMECs following (**D**) normoxia 3h or (**E**) HGD 3h followed by 0, 12, and 24h simulated reperfusion normalized to the housekeeping gene (GAPDH) and expressed as 2^−Δ*Ct*^. n=4-8. (**F**) Line graph depicts summary of the relative temporal expression of *PTGS2*, *IL-1β* and *ICAM1* following either normoxia 3h or HGD 3h followed by 0, 12, and 24h simulated reperfusion normalized to the housekeeping gene (GAPDH) and expressed as 2^−Δ*Ct*^. n=4-8. Comparisons were made using a Two-Way ANOVA with Tukey’s post-hoc test. *P<0.05, **P<0.01, ***P<0.001, ****P<0.0001. (**A**) *CLDN5* (3h/0h) vs. *CLDN5* (3h/12h) ^####^ P < 0.0001, *CLDN5* (3h/0h) vs. *CLDN5* (3h/24h) ^####^ P < 0.0001, *TJP1* (3h/0h) vs. *TJP1* (3h/24h) ^^^^^^ P < 0.0001, *TJP1* (3h/12h) vs. *TJP1* (3h/24h) ^++++^ P < 0.0001. (**A**) *CLDN5* (3h/0h) vs. *CLDN5* (3h/12h) ^####^ P < 0.0001, *CLDN5* (3h/0h) vs. *CLDN5* (3h/24h) ^####^ P < 0.0001, *TJP1* (3h/0h) vs. *TJP1* (3h/12h) ^^^^^^ P < 0.0001, *TJP1* (3h/12h) vs. *TJP1* (3h/24h) ^++++^ P < 0.0001. (**B**) *PTGS2* (3h/0h) vs. *PTGS2* (3h/12h) ^####^ P < 0.0001, *PTGS2* (3h/0h) vs. *PTGS2* (3h/24h) ^####^ P < 0.0001, *ICAM1* (3h/0h) vs. *ICAM1* (3h/12h) ^^^^^^ P < 0.0001, *ICAM1* (3h/0h) vs. *ICAM1* (3h/24h) ^^^^^^ P < 0.0001. (**B**) *PTGS2* (3h/0h) vs. *PTGS2* (3h/24h) ^####^ P < 0.0001, *PTGS2* (3h/12h) vs. *PTGS2* (3h/24h) ^^^^ P = 0.0013.

In addition to evaluating the direct temporal effect on barrier markers, we next sought to evaluate changes within the inflammatory markers PTGS2, IL-1*β*, and ICAM1 at the same timepoints following exposure to either normoxia or HGD/R. In HBMECs exposed to 3h/0h normoxia relatively higher mRNA expression levels of PTGS2 and ICAM1 compared to IL-1*β* were detected (**Fig. 6D**). These results remained relatively consistent at normoxic time points 3h/12h and 3h/24h of apart from slightly decreased ICAM1 expression at 3h/12h. Moreover, we observed an overall decrease in PTGS2 and ICAM1 levels in HBMECs at 3h/12h and 3h/24h compared to levels measured at 3h/0h, potentially suggesting a decrease in basal inflammatory marker transcription overtime (**Fig. 6D**). In contrast, in HBMECs exposed to HGD/R we observed a similar pattern of increased PTGS2 and ICAM1 levels relative to IL-1*β* at all timepoints (**Fig. 6E**). Showcasing a temporal dependent increase in PTGS2 levels at 3h/24h in HBMECs exposed to HGD/R. These findings potentially suggest that of the three markers utilized in this study, the most profound effect of HGD/R was observed to be on increased PTGS2 compared to IL-1*β* and ICAM1 (**Fig. 6E-F**). This was similarly observed in HBMECs exposed to HGD/R (6h/12h) in which PTGS2 and ICAM1 levels were increased (**Supplemental Fig. 3B**).

### 3.7. Correlation of S1PR1-3 with markers of barrier integrity and vascular inflammation

Given the previously established connections between S1PRs and endothelial barrier properties (79-81) as well as cerebral inflammatory responses (82-85) we next sought to address the co-expression of S1PR1-3 with endothelial barrier and inflammatory markers. We first observed that S1PR1 expression in HBMECs was positively correlated with CLDN5 expression and inversely correlated with OCLN, PTGS2, and IL-1β (**Fig. 7A**). In contrast, we observed that HBMEC S1PR2 levels were positively correlated with TJP1, but inversely correlated with CLDN5 (**Fig. 7B**). Interestingly, we did not observe any significant correlations between S1PR2, and the inflammatory markers utilized in this study; however, we did observe a trend of inverse correlation between S1PR2 and ICAM1 (p=0.0641) (**Fig. 7B**). Finally, there was a positive correlation between S1PR3 levels and OCLN as well as IL-1β and a negative correlation with CLDN5 (**Fig. 7C**). Together these data suggest that within HBMECs S1PR1 may be involved in improving barrier integrity via CLDN5 and reducing inflammation via PTGS2 and IL-1β. Additionally, S1PR2 and S1PR3 may be involved in dysregulating endothelial barrier function and exacerbating inflammation (**Supplemental Table 1**). These findings further support that endothelial S1PR1 may be a potential therapeutic target for the mitigation of ischemia reperfusion-like sequalae.

**Figure 7.**
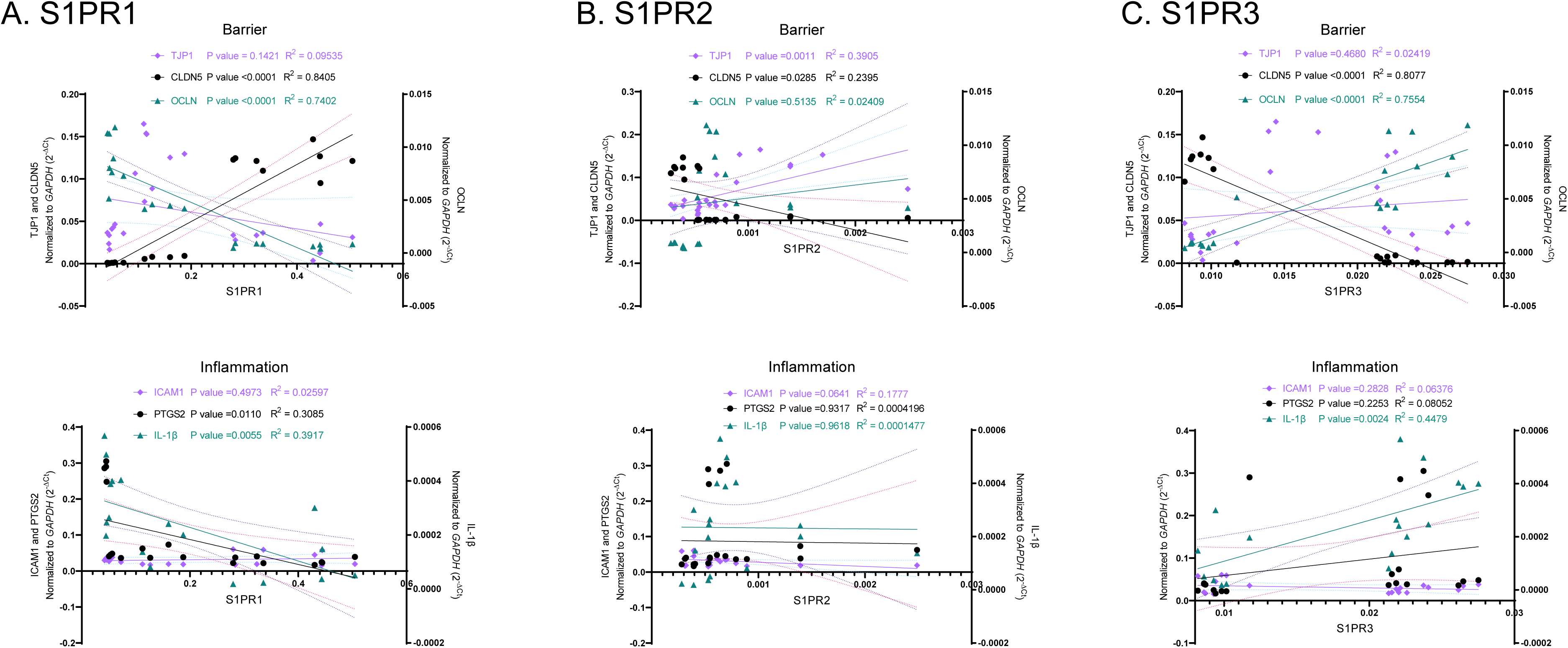
Correlation Between Expression of S1PR 1-3 and Endothelial Barrier and Pro-Inflammatory Mediators. Graphs depicting (**A**) S1PR1, (**B**) S1PR2, (**C**) S1PR3 mRNA expression (x-axis) in HBMECs isolated at HGD 3h followed by simulated reperfusion at 0, 12 and 24h exposure and the corresponding expression of barrier markers including *TJP1*, *CLDN5*, and *OCLN*, as well as inflammatory mediators *ICAM1*, *PTGS2*, and *IL-1β* (y-axis).

### 3.8. Ozanimod, attenuates IL-1*β*-mediated HBMEC MMP-9/2

Based on our observations of HGD/R-induced increases in IL-1β in this study, along with previous findings identifying IL-1β as a key pro-inflammatory cytokine involved in BBB permeability, inflammatory responses, and cell death during AIS pathology(60), we sought to investigate the potential protective effects of S1PR1 activation. Given the established anti-inflammatory role of S1PR1 signaling within the vasculature(86-88), we tested whether ozanimod, a selective S1PR1 agonist, could attenuate the detrimental effects of IL-1β on HBMECs (**Fig. 8A**). Therefore, we first assessed the direct contributions of recombinant IL-1*β* exposure on mRNA markers of HBMEC function, barrier integrity, and inflammation. We observed 24h after administration of recombinant IL-1*β* (400pg/mL) HBMEC S1PR1, ICAM-1, LOX-1, IL-1*β*, and MMP-9, mRNA was increased (**Fig. 8B**). Additionally, we observed a decrease in eNOS mRNA levels (**Fig. 8B**). We also observed in contrast to our hypothesis a concomitant increase in claudin-5 and TIMP-1 mRNA following recombinant IL-1*β* exposure (**Fig. 8B**). These data indicate that the previously observed increase in HBMEC S1PR1 and claudin-5 mRNA following 3h of HGD may be in part mediated by an IL-1*β* mechanism. Moreover, these data suggest that our prior study of observed increases in HBMEC MMP-9 mRNA following 3h of HGD exposure (44) may be in part mediated by an IL-1*β* related cellular mechanism. Given our prior observations regarding the protective effect of ozanimod on HBMECs following HGD, we next investigated the attenuative effects of the selective S1PR1 ligand on recombinant IL-1*β* exposure to further elucidate potential protective mechanisms of S1PR1 within the cerebrovascular endothelium. Ozanimod was given 6h post-administration of recombinant IL-1*β*, to represent a potential therapeutic window of S1PR1 activation following the onset of ischemic injury. Following 18h post-ozanimod delivery and 24h post-recombinant IL-1*β* delivery we observed a decrease in recombinant IL-1*β*-mediated MMP-9 mRNA (**Fig. 8C**). These data suggest that selective S1PR1 activation may be attenuating IL-1*β*-mediated increases in MMP-9 which has been shown to be a key player in the progression of AIS pathogenesis. We further investigated the contributions of recombinant IL-1*β* exposure on MMP-9 as well as MMP-2 activity and observed increased MMP-9 activity following 24h-post administration of recombinant IL-1*β* (**Fig. 8D**). Interestingly, while there was an increase in MMP-9 activity following recombinant IL-1*β* exposure, we did not observe an increase in MMP-2 activity (**Fig. 8D**). These results mirror our prior observations of MMP-9/2 activity within the HBMECs exposed to HGD injury(44). Moreover, we observed that ozanimod, selective S1PR1 ligand, attenuated both MMP-9 and MMP-2 activity (**Fig. 8D**). These observations align with our prior observations that ozanimod via S1PR1 attenuates HBMEC derived MMP-9 and MMP-2 activity following HGD exposure(89). Suggesting that an IL-1*β* mechanism may in part be underlying the observed increase in MMP-9 and that ozanimod, potentially via S1PR1, is attenuating this IL-1*β* mechanism within the HBMECs. In addition to recombinant IL-1*β*, we also utilized cobalt chloride (CoCL_2_; 100*μ*M) to assess the potential contributions of hypoxia inducible factors on HBMEC function, barrier integrity, and inflammation. Similar to our observations of recombinant IL-1*β* exposure, CoCL_2_ exposure induced an increase in LOX-1 and MMP-9 mRNA as well as a decrease in eNOS mRNA (**Supplemental Fig. 4**). However, in contrast to recombinant IL-1*β* exposure, CoCL_2_ exposure decreased ZO-1 mRNA. Additionally, the observed increases in LOX-1 and MMP-9 mRNA following CoCL_2_ exposure was less than that observed following exposure to IL-1*β*. These data together suggest that it may be the inflammatory response that, potentially via an IL-1*β* mechanism that results in increased S1PR1 mRNA rather than a hypoxia mediated mechanism at this time point. Additionally, it may be that IL-1*β* in part underlies the increases in HBMEC MMP-9 mRNA and enzyme activity following HGD exposure and that S1PR1 activation attenuates these increases in part by attenuating IL-1*β*-mediated mechanisms.

**Figure 8.**
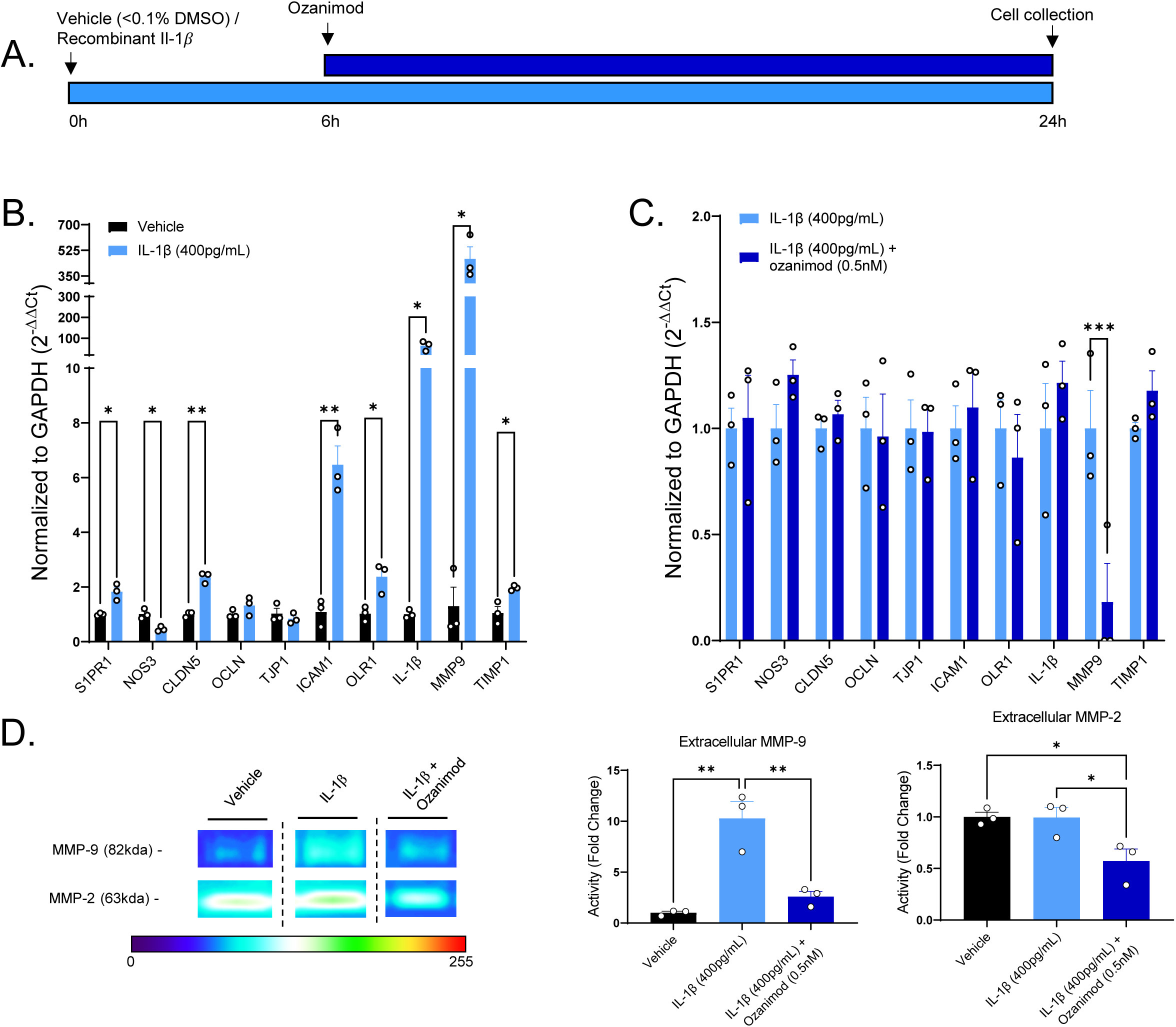
S1PR1, Barrier, and Inflammatory mRNA Expression and Extracellular MMP-9/2 Activity Following Recombinant IL-1β. (**A**) Treatment timeline of HBMECs treated with a single dose of either vehicle or recombinant IL-1β (400pg/mL) given at time 0h, followed by either collection at 24h or administration of ozanimod (0.5nM) at time 6h and collection at time 24h. (**B**) qRT-PCR graph of *S1PR1*, *NOS3*, *CLDN5*, *OCLN*, *TJP1*, *ICAM1*, *OLR1*, *IL-1β*, *MMP9*, and *TIMP1* mRNA expression in HBMECs following vehicle or recombinant IL-1β (400pg/mL) normalized to the housekeeping gene (GAPDH) and expressed as 2^−ΔΔ*Ct*^. n=3. (**C**) qRT-PCR graph of *S1PR1*, *NOS3*, *CLDN5*, *OCLN*, *TJP1*, *ICAM1*, *OLR1*, *IL-1β*, *MMP9*, and *TIMP1* mRNA expression in HBMECs following recombinant IL-1*β* (400pg/mL) exposure alone or treated with ozanimod (0.5nM) normalized to the housekeeping gene (GAPDH) and expressed as 2^−ΔΔ*Ct*^. n=3. (**D**) Representative zymography gel demonstrating the band migration of MMP-9 (82kDa) and MMP-2 (63kDa) in HBMECs following exposure to either vehicle, recombinant IL-1*β* (400pg/mL) exposure, or recombinant IL-1*β* (400pg/mL) exposure and ozanimod (0.5nM). Graphs illustrate quantification of extracellular MMP-9 and MMP-2 enzymatic activity expressed as fold change to vehicle. n=3. Multiple comparisons were performed with one-way ANOVA with Tukey’s post-hoc tests. *P<0.05, **P<0.01.

## 4. DISCUSSION

In this study we have demonstrated that 1) HGD induced an increase in HBMEC S1PR1 and 3 as well as a decrease in HBVSMC S1PR1 and 3; 2) HGD/R injury within HBMECs induces a temporal-dependent decrease in S1PR1 and tight junction marker claudin-5, as well as an increase in pro-inflammatory mediators and a paradoxical increase in occludin mRNA; 3) HBMEC S1PR1 and S1PR2/3 are positively and negatively correlated with claudin-5 expression, respectively; and 4) HBMECs may undergo a phenotypic transformation during the simulated reperfusion phase in which occludin expression predominates over claudin-5. We concomitantly demonstrated the contributions of human recombinant IL-1*β* in dysregulating markers of HBMEC function, barrier integrity, and inflammation as well as the attenuative effects of selective S1PR1 ligand, ozanimod, on attenuating MMP-9/2. These observations coincide with previous studies which show that following ischemic injury, S1PR2 activation leads to detrimental vascular outcomes whereas the activation of S1PR1 elicits beneficial vascular effects. Given this interpretation, we arrived at the concept of a tipping scale, in which a potential therapeutic goal could be to selectively activate the beneficial receptor to override the impact of the detrimental receptor signaling. We believe selective pharmacological modulation of S1PR1, as shown in this study and our previous investigations(44, 45), is needed to tilt the tipping scale in favor of eliciting beneficial cerebrovascular effects through S1PR1 signaling. Therefore, we hypothesize that by administering selective S1PR1 ligands such as ozanimod, S1PR2 activation could be overridden. This hypothesis will require further investigation into the activation of all three receptors following ischemic injury, as well as investigation of the efficacy of selective ligands in their modulation of the receptors.

S1PR1-3 have been reported to be widely expressed in human tissues including the cardiovascular system, nervous system, and immune system(90). On the other hand, S1PR4 is almost exclusively expressed in lymphoid tissue and S1PR5 is highest expressed in the central nervous system(7, 8). However, the relative expression profile of S1PR1-3 at the cerebrovascular level has not been well characterized. It is important to better understand the relative mRNA and protein expression of S1PR1-3 as this may indicate receptor availability for the endogenous S1P ligand and for selective pharmacological ligands and inhibitors that can modulate these receptors. In this study, we found that among the S1PR subtypes, S1PR1 mRNA levels were the highest in HBMECs. (**Figs. 1A & F**). These data suggest that abundant S1PR1 protein may be available for binding of S1P or a selective modulator, leading to beneficial vascular outcomes. Our data highlights that, in HBMECs, S1PR1 mRNA and protein levels increase following 3h HGD exposure (**Figs. 1C & 2A**) which are both relatively decreased at 3h/24h of HGD/R (**Figs. 5A & 5E**), promoting this receptor as a potential early therapeutic target for ischemic injury.

This hypothesis is supported by other studies which have demonstrated that S1PR1 signaling may play a protective role of maintaining barrier function and protecting against vascular injury both *in vitro* and *in vivo*(91-93). Moreover, activation of S1PR1 by a selective S1PR1 modulator, ozanimod, has been shown to attenuate an *in vitro*-ischemic injury-induced reduction in endothelial resistance(10). S1PR1 signals through the G_i_ protein subunit, activating a variety of downstream molecules including PLC, PI3K/Akt, and Ras/ERK, leading to vascular outcomes including angiogenesis, cell survival, proliferation, inflammation, vasorelaxation, and barrier integrity(86-88). With the goal of maintaining a patent and intact vessel following stroke, one of the key mediators in achieving this goal is eNOS, which is the primary generator of NO in endothelial cells. eNOS is activated via the PI3K/Akt pathway, which is integral to the G_i_ signaling cascade through which S1PR1 exclusively functions. One study has previously shown that administration of S1PR1-specific agonists induces the phosphorylation of eNOS in endothelial cells of leptomeningeal collaterals and capillary endothelial cells in the *peri*-infarct area in the early stages of ischemic stroke(92). Beneficial effects of S1PR1 activation through the Akt/eNOS signaling pathway have also been shown in the cardiovascular system by inhibiting the proliferation and migration of fibroblast and transformation of myofibroblast(94). Incorporating the data from this study, we demonstrate the relatively high basal expression of S1PR1 in HBMECs (**Fig. 1F**) and the initial increase in S1PR1 mRNA and protein levels within the HBMECs following HGD injury. We believe this implicates S1PR1 as a therapeutic target following acute ischemic injury. This is further supported by a previous study which also implemented a cell culture model of HBMECs and found that upregulation of S1PR1 protected against oxygen-glucose deprivation and reoxygenation-induced BBB permeability(95). While we observed what seems to be a compensatory mechanism with increased endothelial S1PR1 following HGD exposure, we also observed an interesting paradoxical effect of HGD exposure on S1PR1 mRNA in HBVSMCs. S1PR1 activation has been shown to elicit beneficial effects in HBVSMCs following HGD exposure; our lab previously demonstrated that selective S1PR1 modulation via ozanimod attenuated HGD-induced decreases in cell survival, preserved cytoskeletal integrity, and attenuated vacuolization formation and autophagic flux, thereby contributing to HBVSMC health post-ischemic insult(49, 56). Our data from the current study suggest that S1PR1 may serve as a primary target within the HBMECs rather than HBVSMCs due to the relative higher expression of S1PR2 and S1PR3 in the HBVSMCs both basally and post-HGD injury. Despite this, we still believe that S1PR1 may serve as a stronger therapeutic target following cerebral ischemic injury.

Unlike S1PR1, both S1PR2 and S1PR3 are linked to the G_i_, G_q_ and G_12/13_ subunits, introducing an increasingly complex signaling mechanism. S1PR2 has been shown to predominantly signal through the G12/13 pathway, activating molecules including Rho and ROCK, leading to vasoconstriction, endothelial permeability, and decreased cell proliferation(96). The Rho-ROCK pathway has also been strongly linked with vascular permeability and acute vascular inflammation(97, 98). In this study, we showed that HGD did not alter S1PR2 mRNA and protein levels in HBMECs (**Figs. 1D & 2B**) or HBVSMCs (**Supplemental Fig. 1C**). Given the well-studied vascular outcomes of S1PR2 signaling, we suspect that S1PR2 activation may contribute to detrimental effects following ischemic injury. Previous studies reflect this hypothesis through investigating the impact of S1PR2 gene knockouts on the vasculature. At the whole brain level, Kim et al., 2015 outlined that blockade of S1PR2 either by genetic deletion or pharmacologic inhibition resulted in the reduction of brain infarction, brain edema, neurological deficits, BBB permeability, and neuronal death following the transient (t)MCAO mouse model for stroke(43). More recent studies have also examined the impact of S1PR2 inactivation either by gene knockout or pharmacologic inhibition, finding that S1PR2 inactivation results in higher cerebrovascular integrity and decreased BBB breakdown(41, 99). Moreover, S1PR2 is strongly implicated in cerebrovascular inflammation following ischemic injury. Following ischemia, S1PR2 knockout mice exhibited decreased levels of the inflammatory markers TNF-a and IL-6 as well as reduced microglial activation(84). Our current data describes S1PR2 as relatively more expressed under basal conditions in HBVSMCs than in HBMECs and trended to be higher expressed in HBVSMCs following HGD exposure. Thus, we consider the smooth muscle cellular response to increased S1PR2 expression which has been elucidated through the Rho-ROCK pathway. Tang et al. 2018 demonstrated that RhoA/ROCK signaling induced vascular remodeling and vascular smooth muscle cell phenotypic switching from a quiescent contractile phenotype to a proliferative synthetic phenotype(100). Additionally, it has been previously demonstrated that stimulation of S1PR2 in murine aortic vascular smooth muscle cells induces transcriptional activation of smooth muscle differentiation markers SM22 and smooth muscle *α*-actin, indicators of the contractile phenotype, suggesting that S1PR2 is integral for dedifferentiation of HBVSMCs to the synthetic phenotype(101). These findings complement our lab’s previous work, in which we observed HBVSMC phenotypic switching following HGD exposure with reduced levels of contractile markers including smoothelin, vimentin, myosin light chain kinase (MLCK), and alpha smooth muscle actin (a-SMA)(49). Interestingly, our lab observed that selective modulation of S1PR1 via ozanimod attenuated the HGD-induced phenotypic switching, suggesting that S1PR1 activation may be able to override the S1PR2-induced Rho/ROCK signaling to prevent detrimental vascular cell responses(49).

In contrast to S1PR2, S1PR3 has been shown to primarily work through the G_q_ subunit, signaling to PLC which increases free Ca^2+^ levels, which leads to eNOS activation in endothelial cells to induce vasorelaxation, but concomitantly S1PR3 acting directly on smooth muscle cells leads to vasoconstriction(102). With these varying effects in the two cell types, our findings from this study demonstrating increased endothelial S1PR3 mRNA and decreased smooth muscle S1PR3 mRNA following HGD might suggest a compensatory mechanism at the transcriptional level following exposure to ischemic-like conditions. Similar to the effects of S1PR1 activation, we believe S1PR3 signaling may also contribute to beneficial vascular outcomes post-ischemic insult. Despite this, we believe that S1PR1 may serve as the better therapeutic target because of its exclusive coupling with the G_i_ protein, whereas S1PR3 couples to the G_i_, G_q_ and G_12/13_ proteins and can signal through Rho/ROCK and other pathways that may contribute to poor vascular outcomes.

In the context of ischemia-reperfusion injury (IRI), S1PRs have been shown to exert both beneficial and detrimental effects, depending on the specific receptor subtype and the stage of injury. Numerous studies have supported the claim that S1PR1 activation provides beneficial effects in ischemic injury models. In an *in vivo* study, Brait et al. discovered that the S1PR1-specific agonist LASW1238 significantly reduced infarct size following ischemia/reperfusion of middle cerebral artery(103). Furthermore, in the tMCAO mice model, Hasegawa et al. showed S1PR1-dependent neuronal rescue from apoptosis via AKT and ERK signaling through administration of SEW2871, another S1PR1-specific agonist(93). SEW2871 was also shown to promote lateral branch growth and increase cerebral blood flow in mice with unilateral common carotid artery occlusion (CCAO) surgery(104). These findings provide evidence for S1PR1 activation contributing to the reduction of infarct size and improvement of neurologic function. Interestingly, contrasting effects of S1PR1 action have also been reported. In animal models of tMCAO, inhibition of S1PR1 expression suppressed the proliferation of microglia and reduced the microglia-mediated inflammatory response, ICAM-1 expression, and brain-derived neurotrophic factor expression, thus attenuating BBB dysfunction and brain injury(105). In contrast to S1PR1, S1PR2 signaling has been shown to disrupt cerebrovascular endothelial integrity. Genetic or pharmacological S1PR2 knockout was shown to reduce endothelial MMP-9 expression and attenuate the cerebral edema and hemorrhagic complications after stroke(43). Moreover, inhibition of S1PR2 by a specific antagonist protected the integrity of cerebral endothelial cells and prevented BBB destruction in both *in vitro* and *in vivo* experiments of cerebral ischemic injury(41). In a rat cerebral IRI model and HGD/R model of pre-cerebral cells, there was an upward trend of S1PR2 which was associated with increased BBB permeability presumably via the NF-kB p65 pathway, and that direct targeting of S1PR2 by MiR-149-5p negatively regulated S1PR2, which reduced BBB permeability and improved prognosis of experimental stroke in rats(42). Overall, S1PR2 activity has been linked with detrimental effects in the context of cerebral IRI. S1PR3 activation has been implicated to work similar to S1PR1 following cerebral ischemia, however, its signaling has also been shown to have contrasting impacts which may be due to its coupling to various G protein subunits. One study demonstrated that brain damage during tMCAO IRI in mice was attenuated by S1PR3 regulation of nNOS/NO and oxidative stress(106). Another group found that specific inhibition of S1PR3 by CAY10444 reduced cerebral infarction area and improved neurofunction deficit in tMCAO mice(82). An *in vitro* study showed that phosphorylated FTY720 inhibited the PI3K/NF-kB pathway through S1PR3, which reduced HGD-induced astrocyte injury and the neuroinflammatory response, suggesting a protective effect of S1PR3 in ischemic stroke(107). The contrasting findings of S1PR3 activation in models of IRI warrant further investigation of this receptor subtype and indicate the complexity involved in S1PR3’s linkage to multiple G protein signaling pathways.

In addition to alterations in S1PRs in this study, we observed a temporal dependent alteration in HBMEC tight junction expression following HGD/R exposure from primarily claudin-5 to occludin mRNA. We hypothesize that a transition to primarily expressing occludin instead of claudin-5 in human brain endothelial cells following IRI could indicate a phenotype switch that may have important implications for endothelial barrier integrity and function. Many studies have highlighted the crucial role of both claudin-5 and occludin in maintaining the BBB’s integrity and regulating endothelial barrier function. Claudin-5, considered a "gatekeeper" of neurological function, is vital in restricting paracellular ion movement, contributing to the high electrical resistance of the endothelial barrier(62, 108). Occludin is another significant tight junction protein involved in forming a functional barrier between brain endothelial cells, thus contributing to the overall integrity(109). In the context of IRI, such as that observed in stroke, there might be changes in the expression levels of tight junctions secondary to altered microenvironments and cellular stress. It has been previously shown that occludin degradation can render brain microvascular endothelial cells more vulnerable to reperfusion injury, which suggests that occludin is in part critical for maintaining endothelial cell integrity during injury(110). However, a transition to a state in which the microvascular endothelium primarily express occludin rather than claudin-5 could also indicate a phenotypic switch where the HBMECs prioritize occludin expression to enhance barrier properties and potentially protect against reperfusion injury. In the case of IRI, the HBMECs in this study may experience altered intracellular signaling, leading to changes in the expression of tight junction mRNA and potentially proteins. The upregulation of occludin expression might either be a protective response to reinforce endothelial barrier integrity during this critical period of injury and recovery or a maladaptive response that increases endothelial barrier permeability. Thusly, this transition to primarily expressing occludin instead of claudin-5 in HBMECs following HGD/R injury could signify either a phenotype switch that aims to ultimately enhances or detracts from the BBB’s protective function. Further research is needed to understand the specific molecular mechanisms underlying this transition and its concomitant implications with differential S1PR expression for neurological health and recovery after ischemic events.

We acknowledge that the complex pathophysiological cascade during AIS cannot be exactly modeled in an *in vitro* setting and that HBMECs and HBVSMCs in culture conditions do not exactly mimic the tubular characteristics found *in vivo*. However, *in vitro* studies using human primary cells do allow the investigation of specific basic cellular and molecular mechanisms under conditions of hypoxia or oxygen and glucose deprivation, which reflects what is observed during AIS. Based on our results in this study, we posit that HGD differentially alters S1PR expression in HBVSMCs and HBMECs which suggests that targeting of S1PR1 may be more advantageous in the HBMECs following acute ischemic injury. Moreover, we suggest that HGD/R-induced injury in the HBMECs results in a temporal-dependent concomitant alteration in endothelial barrier phenotype. Additionally, our findings support that S1PR1 within the HBMECs positively correlates with enhanced claudin-5 expression while both S1PR2 and 3 negative correlates with this integral endothelial barrier marker. However, due to the complexity and interconnectivity of GPCRs, it is also likely that stimulation or inactivation of these receptors elicits multiple responses in primary human brain microvascular endothelial cells to attenuate ischemic stroke-like associated pathologies related to endothelial barrier function. Thus, future studies are planned to investigate direct links between S1PRs and the specific mechanism(s) of action in the cerebrovasculature including modulation of Akt, PI3K, Rac1, and upstream mediators in altering endothelial barrier and pro-inflammatory proteins in freshly dissected, uncultured, endothelial cells. Further understanding the role of S1PRs in maintaining the integrity of the cerebrovasculature following an ischemic injury may provide a stronger basis for the selective clinical targeting of S1PR1 with compounds for AIS management and treatment.

## DATA AVAILABILITY

All materials and data will be available from the corresponding author following University of Arizona’s policy of sharing research materials and data.

## DISCLOSURES & DISCLAIMERS

The authors declare no conflict of interest.

## AUTHOR CONTRIBUTIONS

Conceptualization, T.S.W and R.J.G.; methodology, T.S.W, N.B.E, M.C.L, Y.B, and R.J.G.; software, T.S.W, N.B.E and R.J.G.; validation, T.S.W, N.B.E, Y.B, and R.J.G.; formal analysis, T.S.W and R.J.G.; investigation, T.S.W, N.B.E, M.C.L., Y.B, and R.J.G.; resources, R.J.G.; data curation, T.S.W, N.B.E and R.J.G.; writing; review and editing, T.S.W, N.B.E and R.J.G.; visualization, T.S.W and R.J.G.; supervision, R.J.G.; project administration, R.J.G.; funding acquisition, T.S.W and R.J.G. All authors have read and agreed to the published version of the manuscript.

## SUPPLEMENTAL FIGURE LEGENDS

**Supplemental Figure 1.**
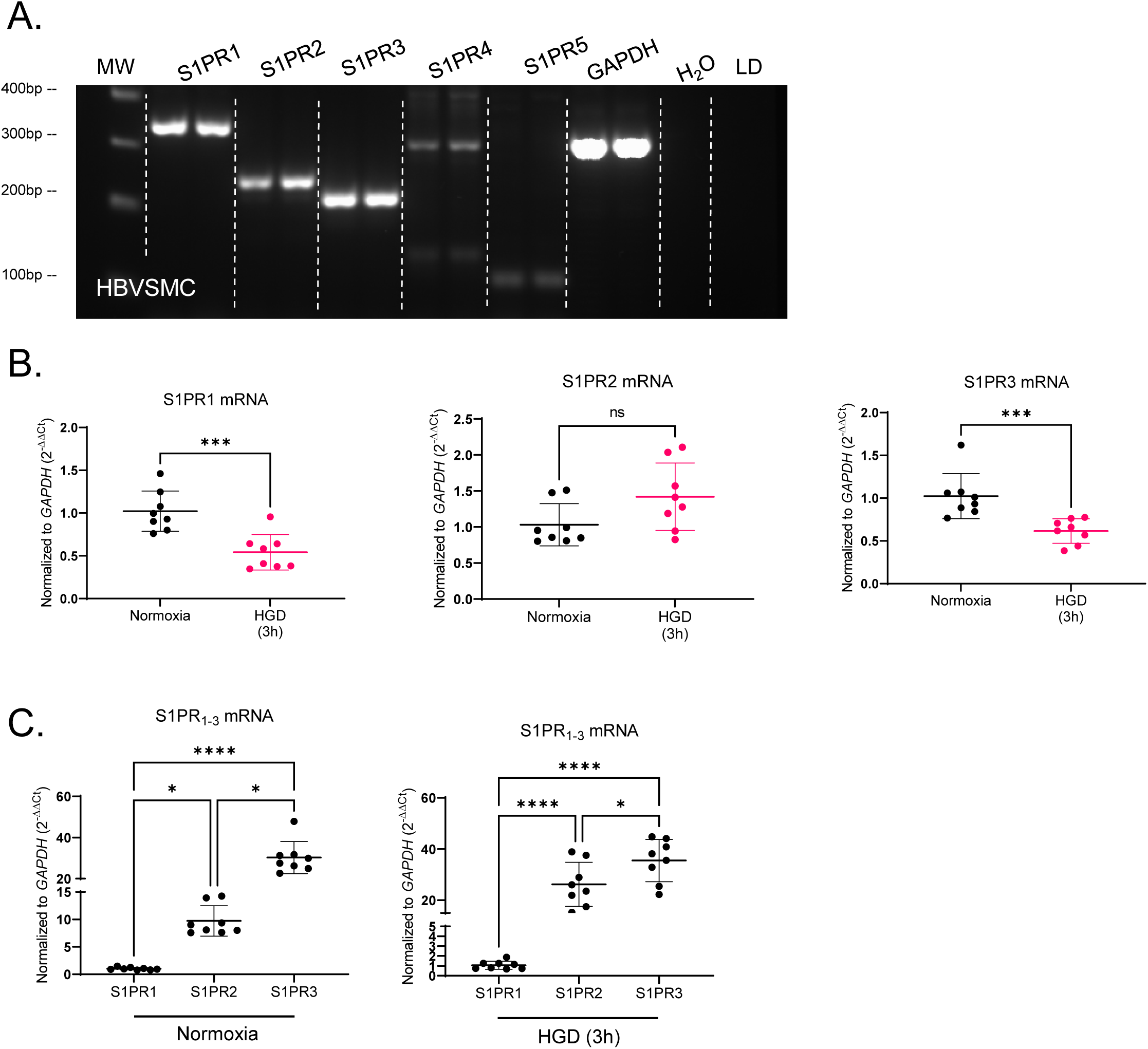
Relative S1PR Types 1-5 mRNA Basal Expression Profile in HBVSMCs Following Normoxia or HGD Exposure. (**A**) RT-PCR ran on 2% agarose gels for sphingosine-1-phosphate receptor (S1PR) types 1-5 to detect receptor mRNA presence in human brain vascular smooth muscle cells (HBVSMCs). GAPDH (loading control) yielded a product size of 309 bp. S1PR1 yielded a product size of 343 bp. S1PR2 yielded a product size of 246 bp. S1PR3 yielded a product size of 215 bp. S1PR4 yielded a product size of 200. S1PR5 yielded a product size of 104 bp. n=2. (**B**) Bar graphs depict qRT-PCR mediated quantitation of S1PR1, S1PR2, and S1PR3 mRNA present within HBVSMCs exposed to either normoxia or HGD (3h) normalized to the housekeeping gene (GAPDH) and expressed as 2^−ΔΔCt^. n=8. (**C**) qRT-PCR graphs of relative S1PR1, S1PR2, S1PR3 mRNA expression in HBVSMCs following normoxia and 3h HGD exposure normalized to the housekeeping gene (GAPDH) and expressed as 2^−ΔΔCt^. n=8. Direct comparisons were performed with unpaired t test with Welch’s correction. Mann-Whitney’s test was utilized to analysis S1PR2 and S1PR3 as these data are non-parametric. *P<0.05, ***P<0.001, ****P<0.0001.

**Supplemental Figure 2.**
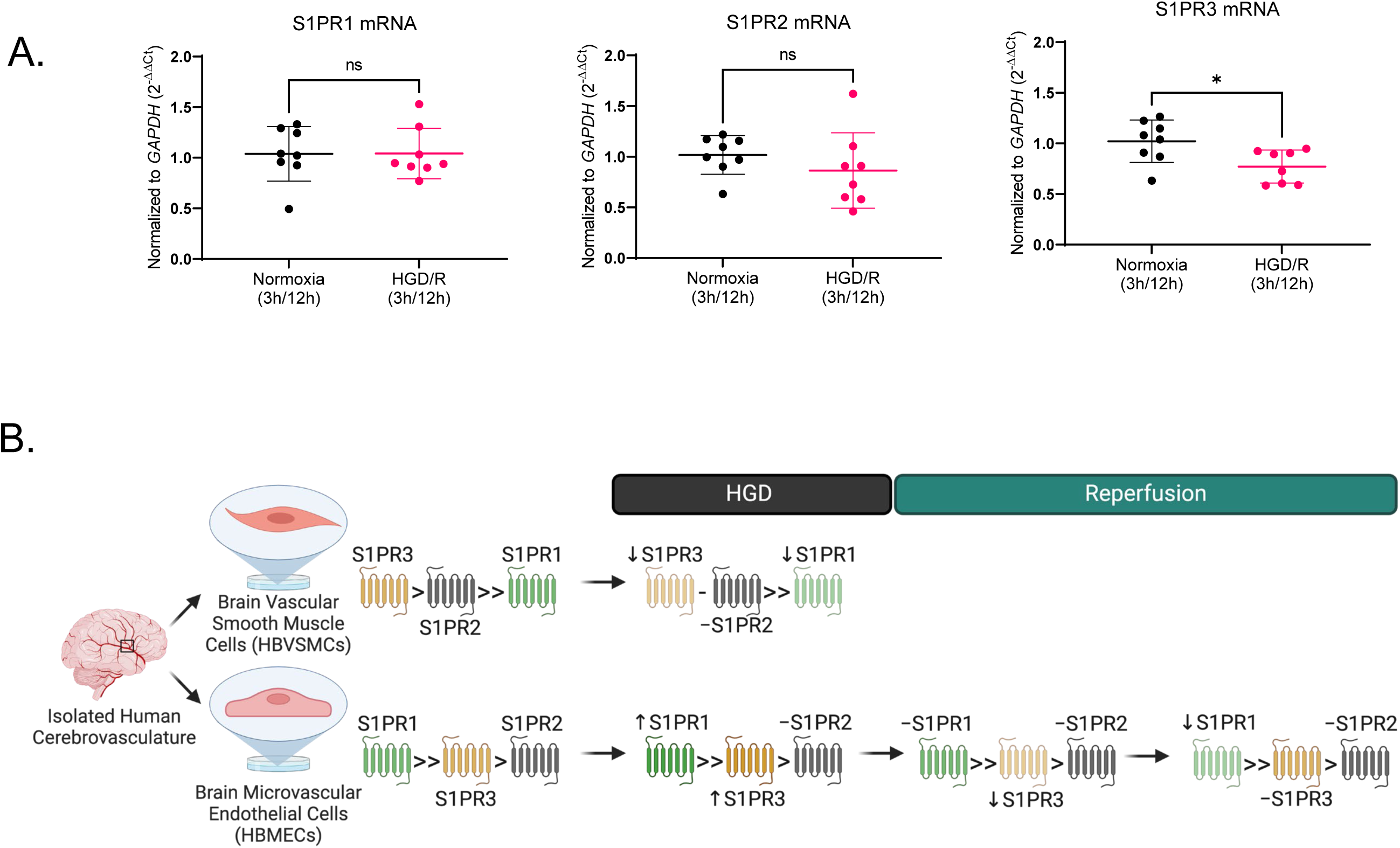
HBMEC S1PR Type 1-3, Inflammatory, and Barrier mRNA Expression Following Hypoxia 3h Plus Glucose Deprivation/Reperfusion 12h (HGD 3h/R 12h) Exposure. qRT-PCR graphs of (**A**) *S1PR1*, *S1PR2*, *S1PR3* mRNA expression in HBMECs following normoxia and HGD 3h /R 12h exposure normalized to the housekeeping gene (GAPDH) and expressed as 2^−ΔΔCt^. n=8. (**B**) Illustrated is a simplistic depiction of the relative expression profile and changes in S1PR types 1-3 in HBVSMCs and HBMECs immediately following 3h of in vitro ischemic-like injury. Additionally, the impact of reperfusion injury on S1PR1-3 within HBMECs is showcased at 12h and 24h of simulated reperfusion. Direct comparisons were performed with unpaired t test with Welch’s correction. Mann-Whitney’s test was utilized to analyze S1PR3 as data are non-parametric. *P<0.05.

**Supplemental 3.**
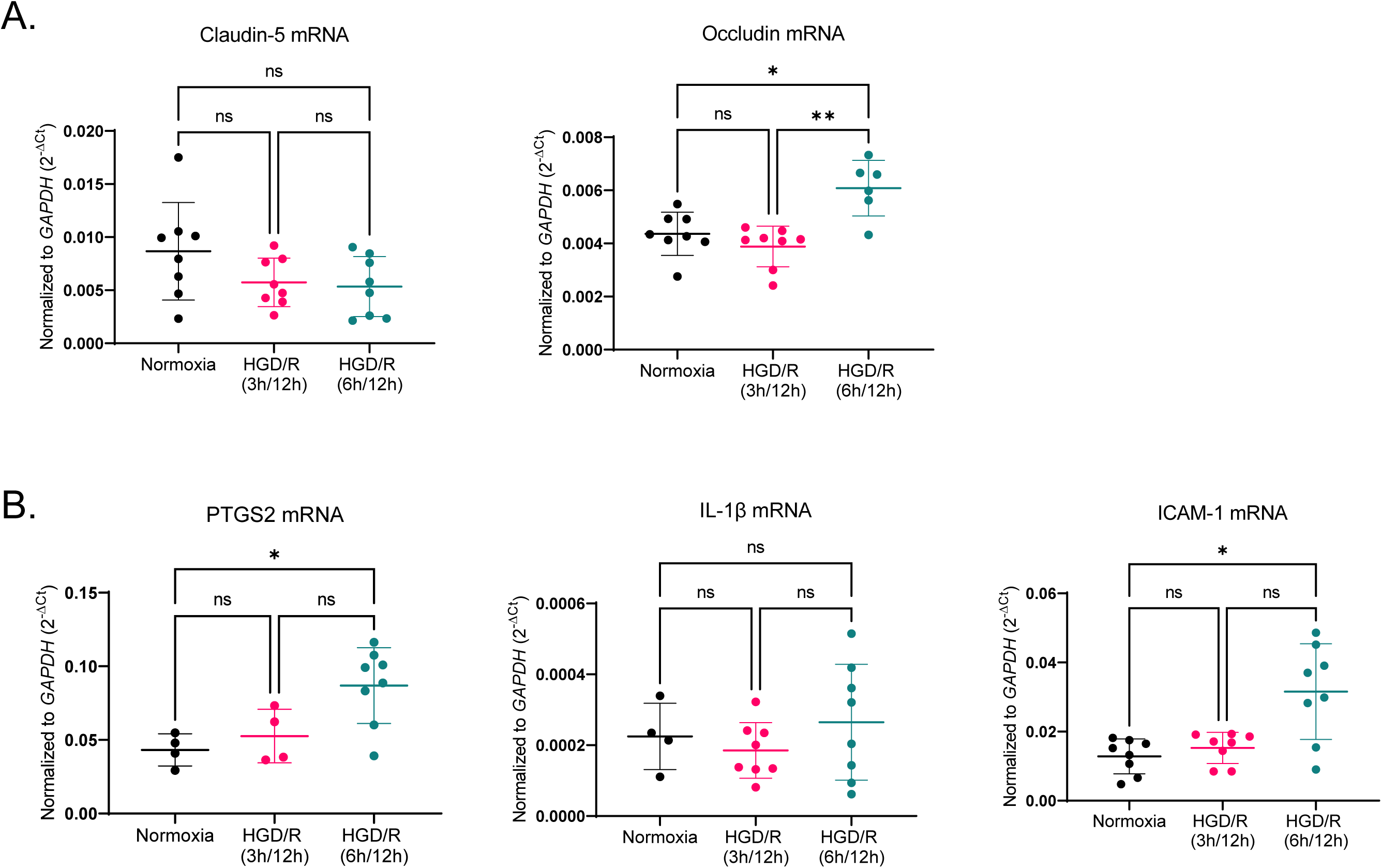
HBMEC Inflammatory and Barrier mRNA Levels Following 3 and 6h Hypoxia Plus Glucose Deprivation with Reperfusion 12h. qRT-PCR graphs of (**A**) *CLDN5* and *OCLN* as well as (**B**) *PTGS2*, *IL-1β*, *ICAM1* mRNA expression in HBMECs following normoxia as well as HGD/R 3h/12h and HGD/R 6h/12h exposure normalized to the housekeeping gene (GAPDH) and expressed as 2^−Δ*Ct*^. Multiple comparisons were performed with one-way ANOVA with Tukey’s post-hoc tests. n=4-8. *P<0.05, **P<0.01.

**Supplemental 4.**
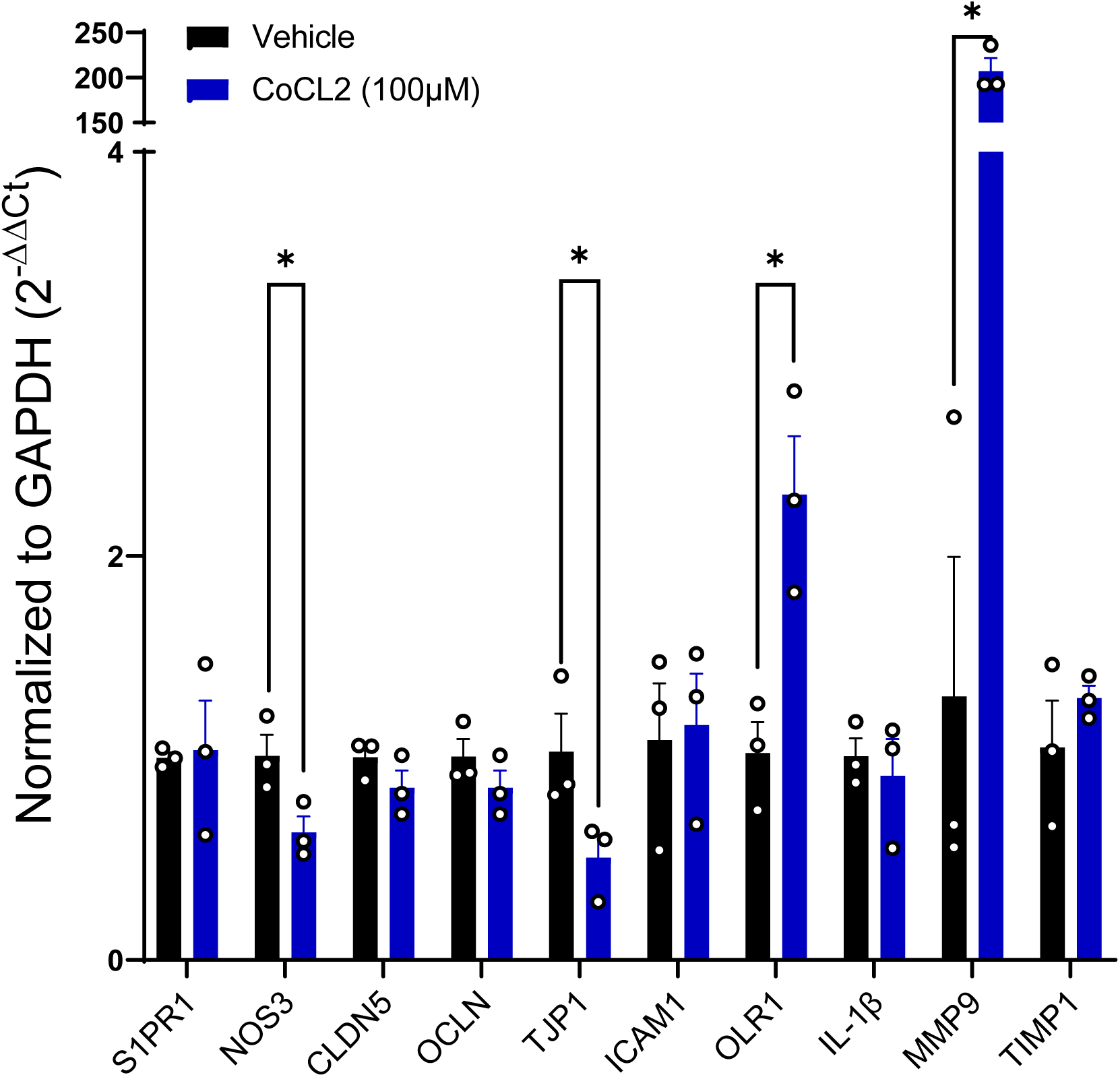
HBMEC S1PR1, Function, Barrier, and Inflammatory mRNA Expression Following CoCL_2_ Exposure. qRT-PCR graph of S1PR1, eNOS, CLDN5, OCLN, TJP1, ICAM1, OLR1, IL-1β, MMP9, and TIMP1 mRNA expression in HBMECs following vehicle or CoCL_2_ (100*μ*M) exposure normalized to the housekeeping gene (GAPDH) and expressed as 2^−ΔΔ*Ct*^. HBMECs were treated with a single dose of either vehicle or CoCL_2_ (100*μ*M) at time 0h followed by collection at 24h. n=3. Direct comparisons were performed with unpaired t test with Welch’s correction. *P<0.05.

## REFERENCES

1. Li X, Stankovic M, Bonder CS, Hahn CN, Parsons M, Pitson SM, et al. Basal and angiopoietin-1-mediated endothelial permeability is regulated by sphingosine kinase-1. Blood. 2008;111(7):3489–97.

2. Camerer E, Regard JB, Cornelissen I, Srinivasan Y, Duong DN, Palmer D, et al. Sphingosine-1-phosphate in the plasma compartment regulates basal and inflammation-induced vascular leak in mice. J Clin Invest. 2009;119(7):1871–9.

3. Obinata H, Hla T. Sphingosine 1-phosphate and inflammation. Int Immunol. 2019;31(9):617–25.

4. Hla T, Maciag T. An abundant transcript induced in differentiating human endothelial cells encodes a polypeptide with structural similarities to G-protein-coupled receptors. J Biol Chem. 1990;265(16):9308–13.

5. Okazaki H, Ishizaka N, Sakurai T, Kurokawa K, Goto K, Kumada M, et al. Molecular cloning of a novel putative G protein-coupled receptor expressed in the cardiovascular system. Biochem Biophys Res Commun. 1993;190(3):1104–9.

6. MacLennan AJ, Browe CS, Gaskin AA, Lado DC, Shaw G. Cloning and characterization of a putative G-protein coupled receptor potentially involved in development. Mol Cell Neurosci. 1994;5(3):201–9.

7. Gräler MH, Bernhardt G, Lipp M. EDG6, a novel G-protein-coupled receptor related to receptors for bioactive lysophospholipids, is specifically expressed in lymphoid tissue. Genomics. 1998;53(2):164–9.

8. Im DS, Heise CE, Ancellin N, O’Dowd BF, Shei GJ, Heavens RP, et al. Characterization of a novel sphingosine 1-phosphate receptor, Edg-8. J Biol Chem. 2000;275(19):14281-6.

9. Prager B, Spampinato SF, Ransohoff RM. Sphingosine 1-phosphate signaling at the blood-brain barrier. Trends Mol Med. 2015;21(6):354–63.

10. Li YJ, Shi SX, Liu Q, Shi FD, Gonzales RJ. Targeted role for sphingosine-1-phosphate receptor 1 in cerebrovascular integrity and inflammation during acute ischemic stroke. Neurosci Lett. 2020;735:135160.

11. Yu L, He L, Gan B, Ti R, Xiao Q, Hu H, et al. Structural insights into sphingosine-1-phosphate receptor activation. Proc Natl Acad Sci U S A. 2022;119(16):e2117716119.

12. Hannun YA, Obeid LM. Principles of bioactive lipid signalling: lessons from sphingolipids. Nat Rev Mol Cell Biol. 2008;9(2):139–50.

13. Rosen H, Goetzl EJ. Sphingosine 1-phosphate and its receptors: an autocrine and paracrine network. Nat Rev Immunol. 2005;5(7):560–70.

14. Pyne S, Pyne NJ. Sphingosine 1-phosphate signalling in mammalian cells. Biochem J. 2000;349(Pt 2):385–402.

15. Garcia JG, Liu F, Verin AD, Birukova A, Dechert MA, Gerthoffer WT, et al. Sphingosine 1-phosphate promotes endothelial cell barrier integrity by Edg-dependent cytoskeletal rearrangement. J Clin Invest. 2001;108(5):689–701.

16. Benechet AP, Menon M, Xu D, Samji T, Maher L, Murooka TT, et al. T cell-intrinsic S1PR1 regulates endogenous effector T-cell egress dynamics from lymph nodes during infection. Proc Natl Acad Sci U S A. 2016;113(8):2182–7.

17. Christensen PM, Liu CH, Swendeman SL, Obinata H, Qvortrup K, Nielsen LB, et al. Impaired endothelial barrier function in apolipoprotein M-deficient mice is dependent on sphingosine-1-phosphate receptor 1. FASEB J. 2016;30(6):2351–9.

18. Oo ML, Chang SH, Thangada S, Wu MT, Rezaul K, Blaho V, et al. Engagement of S1P₁-degradative mechanisms leads to vascular leak in mice. J Clin Invest. 2011;121(6):2290–300.

19. Jung B, Obinata H, Galvani S, Mendelson K, Ding BS, Skoura A, et al. Flow-regulated endothelial S1P receptor-1 signaling sustains vascular development. Dev Cell. 2012;23(3):600–10.

20. Montrose DC, Scherl EJ, Bosworth BP, Zhou XK, Jung B, Dannenberg AJ, et al. S1P₁ localizes to the colonic vasculature in ulcerative colitis and maintains blood vessel integrity. J Lipid Res. 2013;54(3):843–51.

21. Igarashi J, Bernier SG, Michel T. Sphingosine 1-phosphate and activation of endothelial nitric-oxide synthase. differential regulation of Akt and MAP kinase pathways by EDG and bradykinin receptors in vascular endothelial cells. J Biol Chem. 2001;276(15):12420–6.

22. Morales-Ruiz M, Lee MJ, Zöllner S, Gratton JP, Scotland R, Shiojima I, et al. Sphingosine 1-phosphate activates Akt, nitric oxide production, and chemotaxis through a Gi protein/phosphoinositide 3-kinase pathway in endothelial cells. J Biol Chem. 2001;276(22):19672–7.

23. Ishii I, Fukushima N, Ye X, Chun J. Lysophospholipid receptors: signaling and biology. Annu Rev Biochem. 2004;73:321–54.

24. Nofer JR, van der Giet M, Tölle M, Wolinska I, von Wnuck Lipinski K, Baba HA, et al. HDL induces NO-dependent vasorelaxation via the lysophospholipid receptor S1P3. J Clin Invest. 2004;113(4):569–81.

25. Katunaric B, SenthilKumar G, Schulz ME, De Oliveira N, Freed JK. S1P (Sphingosine-1-Phosphate)-Induced Vasodilation in Human Resistance Arterioles During Health and Disease. Hypertension. 2022;79(10):2250–61.

26. Salomone S, Potts EM, Tyndall S, Ip PC, Chun J, Brinkmann V, et al. Analysis of sphingosine 1-phosphate receptors involved in constriction of isolated cerebral arteries with receptor null mice and pharmacological tools. Br J Pharmacol. 2008;153(1):140–7.

27. Cantalupo A, Gargiulo A, Dautaj E, Liu C, Zhang Y, Hla T, et al. S1PR1 (Sphingosine-1-Phosphate Receptor 1) Signaling Regulates Blood Flow and Pressure. Hypertension. 2017;70(2):426–34.

28. Lorenz JN, Arend LJ, Robitz R, Paul RJ, MacLennan AJ. Vascular dysfunction in S1P2 sphingosine 1-phosphate receptor knockout mice. Am J Physiol Regul Integr Comp Physiol. 2007;292(1):R440–6.

29. Szczepaniak WS, Pitt BR, McVerry BJ. S1P2 receptor-dependent Rho-kinase activation mediates vasoconstriction in the murine pulmonary circulation induced by sphingosine 1-phosphate. Am J Physiol Lung Cell Mol Physiol. 2010;299(1):L137–45.

30. Salomone S, Yoshimura S, Reuter U, Foley M, Thomas SS, Moskowitz MA, et al. S1P3 receptors mediate the potent constriction of cerebral arteries by sphingosine-1-phosphate. Eur J Pharmacol. 2003;469(1-3):125–34.

31. Takuwa Y, Du W, Qi X, Okamoto Y, Takuwa N, Yoshioka K. Roles of sphingosine-1-phosphate signaling in angiogenesis. World J Biol Chem. 2010;1(10):298–306.

32. Watanabe S, Usui-Kawanishi F, Karasawa T, Kimura H, Kamata R, Komada T, et al. Glucose regulates hypoxia-induced NLRP3 inflammasome activation in macrophages. J Cell Physiol. 2020;235(10):7554–66.

33. Zhang W, Smith C, Howlett C, Stanimirovic D. Inflammatory activation of human brain endothelial cells by hypoxic astrocytes in vitro is mediated by IL-1beta. J Cereb Blood Flow Metab. 2000;20(6):967–78.

34. Boutin H, LeFeuvre RA, Horai R, Asano M, Iwakura Y, Rothwell NJ. Role of IL-1alpha and IL-1beta in ischemic brain damage. J Neurosci. 2001;21(15):5528–34.

35. Ghezzi P, Dinarello CA, Bianchi M, Rosandich ME, Repine JE, White CW. Hypoxia increases production of interleukin-1 and tumor necrosis factor by human mononuclear cells. Cytokine. 1991;3(3):189–94.

36. Deb P, Sharma S, Hassan KM. Pathophysiologic mechanisms of acute ischemic stroke: An overview with emphasis on therapeutic significance beyond thrombolysis. Pathophysiology. 2010;17(3):197–218.

37. Maceyka M, Harikumar KB, Milstien S, Spiegel S. Sphingosine-1-phosphate signaling and its role in disease. Trends Cell Biol. 2012;22(1):50–60.

38. Jozefczuk E, Guzik TJ, Siedlinski M. Significance of sphingosine-1-phosphate in cardiovascular physiology and pathology. Pharmacol Res. 2020;156:104793.

39. Salas-Perdomo A, Miró-Mur F, Gallizioli M, Brait VH, Justicia C, Meissner A, et al. Role of the S1P pathway and inhibition by fingolimod in preventing hemorrhagic transformation after stroke. Sci Rep. 2019;9(1):8309.

40. Gaengel K, Niaudet C, Hagikura K, Laviña B, Siemsen BL, Muhl L, et al. The sphingosine-1-phosphate receptor S1PR1 restricts sprouting angiogenesis by regulating the interplay between VE-cadherin and VEGFR2. Dev Cell. 2012;23(3):587–99.

41. Cao C, Dai L, Mu J, Wang X, Hong Y, Zhu C, et al. S1PR2 antagonist alleviates oxidative stress-enhanced brain endothelial permeability by attenuating p38 and Erk1/2-dependent cPLA. Cell Signal. 2019;53:151–61.

42. Wan Y, Jin HJ, Zhu YY, Fang Z, Mao L, He Q, et al. MicroRNA-149-5p regulates blood-brain barrier permeability after transient middle cerebral artery occlusion in rats by targeting S1PR2 of pericytes. FASEB J. 2018;32(6):3133–48.

43. Kim GS, Yang L, Zhang G, Zhao H, Selim M, McCullough LD, et al. Critical role of sphingosine-1-phosphate receptor-2 in the disruption of cerebrovascular integrity in experimental stroke. Nat Commun. 2015;6:7893.

44. Wendt TS, Gonzales RJ. Ozanimod Differentially Preserves Human Cerebrovascular Endothelial Barrier Proteins and Attenuates MMP-9 Activity Following In Vitro Acute Ischemic Injury. Am J Physiol Cell Physiol. 2023.

45. Wendt TS, Li YJ, Gonzales RJ. Ozanimod, an S1PR1 ligand, attenuates hypoxia plus glucose deprivation-induced autophagic flux and phenotypic switching in human brain VSM cells. Am J Physiol Cell Physiol. 2021;320(6):C1055–C73.

46. O’Carroll SJ, Kho DT, Wiltshire R, Nelson V, Rotimi O, Johnson R, et al. Pro-inflammatory TNFα and IL-1β differentially regulate the inflammatory phenotype of brain microvascular endothelial cells. J Neuroinflammation. 2015;12:131.

47. Bayat M, Razavi Moosavi N, Karimi N, Rahimi M, Borhani-Haghighi A. Increased Serum Levels of IL-1β after Ischemic Stroke are Inversely Associated with Vitamin D. Iran J Immunol. 2023;20(3):359–67.

48. Page S, Munsell A, Al-Ahmad AJ. Cerebral hypoxia/ischemia selectively disrupts tight junctions complexes in stem cell-derived human brain microvascular endothelial cells. Fluids Barriers CNS. 2016;13(1):16.

49. Wendt TS, Li YJ, Gonzales RJ. Ozanimod, an S1PR1 ligand, attenuates hypoxia plus glucose deprivation induced autophagic flux and phenotypic switching in human brain VSM cells. Am J Physiol Cell Physiol. 2021.

50. Du J, Zeng C, Li Q, Chen B, Liu H, Huang X, et al. LPS and TNF-α induce expression of sphingosine-1-phosphate receptor-2 in human microvascular endothelial cells. Pathol Res Pract. 2012;208(2):82–8.

51. Panetti TS. Differential effects of sphingosine 1-phosphate and lysophosphatidic acid on endothelial cells. Biochim Biophys Acta. 2002;1582(1-3):190–6.

52. Straub AC, Klei LR, Stolz DB, Barchowsky A. Arsenic requires sphingosine-1-phosphate type 1 receptors to induce angiogenic genes and endothelial cell remodeling. Am J Pathol. 2009;174(5):1949–58.

53. Bolick DT, Srinivasan S, Kim KW, Hatley ME, Clemens JJ, Whetzel A, et al. Sphingosine-1-phosphate prevents tumor necrosis factor-{alpha}-mediated monocyte adhesion to aortic endothelium in mice. Arterioscler Thromb Vasc Biol. 2005;25(5):976–81.

54. Lee MJ, Thangada S, Claffey KP, Ancellin N, Liu CH, Kluk M, et al. Vascular endothelial cell adherens junction assembly and morphogenesis induced by sphingosine-1-phosphate. Cell. 1999;99(3):301–12.

55. Duru EA, Fu Y, Davies MG. Role of S-1-P receptors and human vascular smooth muscle cell migration in diabetes and metabolic syndrome. J Surg Res. 2012;177(2):e75–82.

56. Wendt TS, Li YJ, Gonzales RJ. Ozanimod, an S1PR. Am J Physiol Cell Physiol. 2021;320(6):C1055–C73.

57. Andjelkovic AV, Xiang J, Stamatovic SM, Hua Y, Xi G, Wang MM, et al. Endothelial Targets in Stroke: Translating Animal Models to Human. Arterioscler Thromb Vasc Biol. 2019;39(11):2240–7.

58. Iadecola C, Gorelick PB. The Janus face of cyclooxygenase-2 in ischemic stroke: shifting toward downstream targets. Stroke. 2005;36(2):182–5.

59. Wong R, Lénárt N, Hill L, Toms L, Coutts G, Martinecz B, et al. Interleukin-1 mediates ischaemic brain injury via distinct actions on endothelial cells and cholinergic neurons. Brain Behav Immun. 2019;76:126–38.

60. Zhu H, Hu S, Li Y, Sun Y, Xiong X, Hu X, et al. Interleukins and Ischemic Stroke. Front Immunol. 2022;13:828447.

61. Lindsberg PJ, Carpén O, Paetau A, Karjalainen-Lindsberg ML, Kaste M. Endothelial ICAM-1 expression associated with inflammatory cell response in human ischemic stroke. Circulation. 1996;94(5):939–45.

62. Greene C, Hanley N, Campbell M. Claudin-5: gatekeeper of neurological function. Fluids Barriers CNS. 2019;16(1):3.

63. Yuan S, Liu KJ, Qi Z. Occludin regulation of blood-brain barrier and potential therapeutic target in ischemic stroke. Brain Circ. 2020;6(3):152–62.

64. Abdullahi W, Tripathi D, Ronaldson PT. Blood-brain barrier dysfunction in ischemic stroke: targeting tight junctions and transporters for vascular protection. Am J Physiol Cell Physiol. 2018;315(3):C343–C56.

65. Pan R, Yu K, Weatherwax T, Zheng H, Liu W, Liu KJ. Blood Occludin Level as a Potential Biomarker for Early Blood Brain Barrier Damage Following Ischemic Stroke. Sci Rep. 2017;7:40331.

66. Jiao H, Wang Z, Liu Y, Wang P, Xue Y. Specific role of tight junction proteins claudin-5, occludin, and ZO-1 of the blood-brain barrier in a focal cerebral ischemic insult. J Mol Neurosci. 2011;44(2):130–9.

67. Tran JQ, Zhang P, Ghosh A, Liu L, Syto M, Wang X, et al. Single-Dose Pharmacokinetics of Ozanimod and its Major Active Metabolites Alone and in Combination with Gemfibrozil, Itraconazole, or Rifampin in Healthy Subjects: A Randomized, Parallel-Group, Open-Label Study. Adv Ther. 2020;37(10):4381–95.

68. Sugahara K, Maeda Y, Shimano K, Mogami A, Kataoka H, Ogawa K, et al. Amiselimod, a novel sphingosine 1-phosphate receptor-1 modulator, has potent therapeutic efficacy for autoimmune diseases, with low bradycardia risk. Br J Pharmacol. 2017;174(1):15–27.

69. Brossard P, Derendorf H, Xu J, Maatouk H, Halabi A, Dingemanse J. Pharmacokinetics and pharmacodynamics of ponesimod, a selective S1P1 receptor modulator, in the first-in-human study. Br J Clin Pharmacol. 2013;76(6):888–96.

70. Glaenzel U, Jin Y, Nufer R, Li W, Schroer K, Adam-Stitah S, et al. Metabolism and Disposition of Siponimod, a Novel Selective S1P. Drug Metab Dispos. 2018;46(7):1001–13.

71. Saren G, Wong A, Lu YB, Baciu C, Zhou W, Zamel R, et al. Ischemia-Reperfusion Injury in a Simulated Lung Transplant Setting Differentially Regulates Transcriptomic Profiles between Human Lung Endothelial and Epithelial Cells. Cells. 2021;10(10).

72. Sebastião MJ, Gomes-Alves P, Reis I, Sanchez B, Palacios I, Serra M, et al. Bioreactor-based 3D human myocardial ischemia/reperfusion in vitro model: a novel tool to unveil key paracrine factors upon acute myocardial infarction. Transl Res. 2020;215:57–74.

73. Makkos A, Szántai Á, Pálóczi J, Pipis J, Kiss B, Poggi P, et al. A Comorbidity Model of Myocardial Ischemia/Reperfusion Injury and Hypercholesterolemia in Rat Cardiac Myocyte Cultures. Front Physiol. 2019;10:1564.

74. Kulek AR, Undyala VVR, Anzell AR, Raghunayakula S, MacMillan-Crow LA, Sanderson TH, et al. Does Disruption of Optic Atrophy-1 (OPA1) Contribute to Cell Death in HL-1 Cardiomyocytes Subjected to Lethal Ischemia-Reperfusion Injury? Cells. 2022;11(19).

75. Chen L, Luo W, Zhang W, Chu H, Wang J, Dai X, et al. circDLPAG4/HECTD1 mediates ischaemia/reperfusion injury in endothelial cells via ER stress. RNA Biol. 2020;17(2):240–53.

76. Xie X, Zhu T, Chen L, Ding S, Chu H, Wang J, et al. MCPIP1-induced autophagy mediates ischemia/reperfusion injury in endothelial cells via HMGB1 and CaSR. Sci Rep. 2018;8(1):1735.

77. Arkelius K, Wendt TS, Andersson H, Arnou A, Gottschalk M, Gonzales RJ, et al. LOX-1 and MMP-9 Inhibition Attenuates the Detrimental Effects of Delayed rt-PA Therapy and Improves Outcomes After Acute Ischemic Stroke. Circ Res. 2024;134(8):954–69.

78. L L, X W, Z Y. Ischemia-reperfusion Injury in the Brain: Mechanisms and Potential Therapeutic Strategies. Biochem Pharmacol (Los Angel). 2016;5(4).

79. Yanagida K, Liu CH, Faraco G, Galvani S, Smith HK, Burg N, et al. Size-selective opening of the blood-brain barrier by targeting endothelial sphingosine 1-phosphate receptor 1. Proc Natl Acad Sci U S A. 2017;114(17):4531–6.

80. Xiang P, Chew WS, Seow WL, Lam BWS, Ong WY, Herr DR. The S1P. Neurochem Int. 2021;146:105018.

81. Fan X, Chen H, Xu C, Wang Y, Yin P, Li M, et al. S1PR3, as a Core Protein Related to Ischemic Stroke, is Involved in the Regulation of Blood-Brain Barrier Damage. Front Pharmacol. 2022;13:834948.

82. Gaire BP, Song MR, Choi JW. Sphingosine 1-phosphate receptor subtype 3 (S1P. J Neuroinflammation. 2018;15(1):284.

83. Gaire BP, Bae YJ, Choi JW. S1P. Biomol Ther (Seoul). 2019;27(6):522–9.

84. Sapkota A, Gaire BP, Kang MG, Choi JW. S1P. Sci Rep. 2019;9(1):12106.

85. Gaire BP, Choi JW. Sphingosine 1-Phosphate Receptors in Cerebral Ischemia. Neuromolecular Med. 2021;23(1):211–23.

86. Lee MJ, Thangada S, Paik JH, Sapkota GP, Ancellin N, Chae SS, et al. Akt-mediated phosphorylation of the G protein-coupled receptor EDG-1 is required for endothelial cell chemotaxis. Mol Cell. 2001;8(3):693–704.

87. Mehta D, Konstantoulaki M, Ahmmed GU, Malik AB. Sphingosine 1-phosphate-induced mobilization of intracellular Ca2+ mediates rac activation and adherens junction assembly in endothelial cells. J Biol Chem. 2005;280(17):17320–8.

88. Li Q, Chen B, Zeng C, Fan A, Yuan Y, Guo X, et al. Differential activation of receptors and signal pathways upon stimulation by different doses of sphingosine-1-phosphate in endothelial cells. Exp Physiol. 2015;100(1):95–107.

89. Samuel XS, Yu-Jing L, Trevor SW, Kaibin S, Weina J, Qiang L, et al. Selective S1PR1 activation improves brain infarction, neurological outcome, and cerebrovascular endothelial health following experimental ischemic injury. bioRxiv. 2023:2023.03.27.534410.

90. Lu S, She M, Zeng Q, Yi G, Zhang J. Sphingosine 1-phosphate and its receptors in ischemia. Clin Chim Acta. 2021;521:25–33.

91. Burg N, Swendeman S, Worgall S, Hla T, Salmon JE. Sphingosine 1-Phosphate Receptor 1 Signaling Maintains Endothelial Cell Barrier Function and Protects Against Immune Complex-Induced Vascular Injury. Arthritis Rheumatol. 2018;70(11):1879–89.

92. Iwasawa E, Ishibashi S, Suzuki M, Li F, Ichijo M, Miki K, et al. Sphingosine-1-Phosphate Receptor 1 Activation Enhances Leptomeningeal Collateral Development and Improves Outcome after Stroke in Mice. J Stroke Cerebrovasc Dis. 2018;27(5):1237–51.

93. Hasegawa Y, Suzuki H, Sozen T, Rolland W, Zhang JH. Activation of sphingosine 1-phosphate receptor-1 by FTY720 is neuroprotective after ischemic stroke in rats. Stroke. 2010;41(2):368–74.

94. Liu X, Wu J, Zhu C, Liu J, Chen X, Zhuang T, et al. Endothelial S1pr1 regulates pressure overload-induced cardiac remodelling through AKT-eNOS pathway. J Cell Mol Med. 2020;24(2):2013–26.

95. Lin L, Wang Q, Qian K, Cao Z, Xiao J, Wang X, et al. bFGF Protects Against Oxygen Glucose Deprivation/Reoxygenation-Induced Endothelial Monolayer Permeability via S1PR1-Dependent Mechanisms. Mol Neurobiol. 2018;55(4):3131–42.

96. Sanchez T, Skoura A, Wu MT, Casserly B, Harrington EO, Hla T. Induction of vascular permeability by the sphingosine-1-phosphate receptor-2 (S1P2R) and its downstream effectors ROCK and PTEN. Arterioscler Thromb Vasc Biol. 2007;27(6):1312–8.

97. Laufs U, Liao JK. Post-transcriptional regulation of endothelial nitric oxide synthase mRNA stability by Rho GTPase. J Biol Chem. 1998;273(37):24266–71.

98. Zhang G, Yang L, Kim GS, Ryan K, Lu S, O’Donnell RK, et al. Critical role of sphingosine-1-phosphate receptor 2 (S1PR2) in acute vascular inflammation. Blood. 2013;122(3):443–55.

99. Seyedsadr MS, Weinmann O, Amorim A, Ineichen BV, Egger M, Mirnajafi-Zadeh J, et al. Inactivation of sphingosine-1-phosphate receptor 2 (S1PR2) decreases demyelination and enhances remyelination in animal models of multiple sclerosis. Neurobiol Dis. 2019;124:189–201.

100. Tang L, Dai F, Liu Y, Yu X, Huang C, Wang Y, et al. RhoA/ROCK signaling regulates smooth muscle phenotypic modulation and vascular remodeling via the JNK pathway and vimentin cytoskeleton. Pharmacol Res. 2018;133:201–12.

101. Medlin MD, Staus DP, Dubash AD, Taylor JM, Mack CP. Sphingosine 1-phosphate receptor 2 signals through leukemia-associated RhoGEF (LARG), to promote smooth muscle cell differentiation. Arterioscler Thromb Vasc Biol. 2010;30(9):1779–86.

102. An S, Bleu T, Zheng Y. Transduction of intracellular calcium signals through G protein-mediated activation of phospholipase C by recombinant sphingosine 1-phosphate receptors. Mol Pharmacol. 1999;55(5):787–94.

103. Brait VH, Tarrasón G, Gavaldà A, Godessart N, Planas AM. Selective Sphingosine 1-Phosphate Receptor 1 Agonist Is Protective Against Ischemia/Reperfusion in Mice. Stroke. 2016;47(12):3053–6.

104. Ichijo M, Ishibashi S, Li F, Yui D, Miki K, Mizusawa H, et al. Sphingosine-1-Phosphate Receptor-1 Selective Agonist Enhances Collateral Growth and Protects against Subsequent Stroke. PLoS One. 2015;10(9):e0138029.

105. Gaire BP, Lee CH, Sapkota A, Lee SY, Chun J, Cho HJ, et al. Identification of Sphingosine 1-Phosphate Receptor Subtype 1 (S1P. Mol Neurobiol. 2018;55(3):2320–32.

106. Fan X, Chen H, Xu C, Wang Y, Yin P, Li M, et al. Inhibiting Sphingosine 1-Phosphate Receptor Subtype 3 Attenuates Brain Damage During Ischemia-Reperfusion Injury by Regulating nNOS/NO and Oxidative Stress. Front Neurosci. 2022;16:838621.

107. Dong YF, Guo RB, Ji J, Cao LL, Zhang L, Chen ZZ, et al. S1PR3 is essential for phosphorylated fingolimod to protect astrocytes against oxygen-glucose deprivation-induced neuroinflammation via inhibiting TLR2/4-NFκB signalling. J Cell Mol Med. 2018;22(6):3159–66.

108. Zhong X, Luo C, Deng M, Zhao M. Scutellarin-treated exosomes increase claudin 5, occludin and ZO1 expression in rat brain microvascular endothelial cells. Exp Ther Med. 2019;18(1):33–40.

109. Keil JM, Liu X, Antonetti DA. Glucocorticoid induction of occludin expression and endothelial barrier requires transcription factor p54 NONO. Invest Ophthalmol Vis Sci. 2013;54(6):4007–15.

110. Zhang Y, Li X, Qiao S, Yang D, Li Z, Xu J, et al. Occludin degradation makes brain microvascular endothelial cells more vulnerable to reperfusion injury in vitro. J Neurochem. 2021;156(3):352–66.

